# A model of calcium-induced calcium release from the sarcoplasmic reticulum in the smooth muscle cell and its investigation by mathematical modeling

**DOI:** 10.1101/2023.08.12.553083

**Authors:** P. F. Zhuk, S. O. Karakhim, S. O. Kosterin

**Author notes:** Corresponding author: S. O. Karakhim.

## Abstract

A model of calcium-induced calcium release (CICR) from the sarcoplasmic reticulum was developed, simulating the behavior of the smooth muscle cell under conditions of its agonist stimulation. The mathematical model is a system of thirteen differential equations. In the initial (basal) state, the parameters of active and passive transport of Ca^2+^ through both the plasma membrane and the sarcoplasmic reticulum membrane are adjusted.

A study of the model showed that, depending on the model parameters, the cell system can demonstrate two types of calcium concentration changes in the cytosol: a single Ca^2+^ transient and an oscillatory mode.

After stimulation is completed, the cell system returns to the basal state (under *in vivo* conditions) or goes to a new steady-state level (under *in vitro* conditions), except when the cell system is in oscillatory mode. It is shown that the sarcoplasmic reticulum can act both as a passive participant in the process of Ca^2+^ accumulation in the smooth muscle cell, acting as a buffer, and play a major role in this process by significantly increasing the Ca^2+^ concentration in the cytosol, which is initiated by Ca^2+^ entry from the extracellular space.

It was found that after stimulation of the smooth muscle cell, the net Ca^2+^ flux from the cytosol increases due to an increase in Ca^2+^ concentration in the cytosol, while the net Ca^2+^ flux into the cytosol first increases due to an increase in the number of open calcium channels located on the membrane of the sarcoplasmic reticulum. It then begins to decrease due to a decrease in the Ca^2+^ concentration gradient across the membrane of the sarcoplasmic reticulum. Therefore, at a certain time point these fluxes become equal and the process of Ca^2+^ accumulation in the cytosol is terminated. Thus, calcium-induced calcium release is terminated spontaneously, resulting in the formation of a single cytosolic Ca^2+^ transient. As a result of calcium-induced calcium release, the sarcoplasmic reticulum is not completely emptied, but retains quite significant amounts of Ca^2+^.

The possibility of Ca^2+^ redistribution between the three compartments (extracellular space, cytosol and sarcoplasmic reticulum) creates the possibility of oscillation of cytosolic Ca^2+^ concentration. The oscillation amplitude and frequency can remain practically unchanged for a considerable period.

The developed model qualitatively reproduces the results of experimental studies conducted to identify store-operated calcium channels using the inhibitors of the calcium pump of the sarcoplasmic reticulum in a calcium-free medium.

## 1 Introduction

Intracellular free Ca^2+^ is the most important factor in the muscle contraction-relaxation mechanism. It initiates several interrelated reactions involving, in particular, actin, myosin, and ATP, which results in muscle contraction [1-6]. The degree of contraction is determined by the intracellular Ca^2+^ concentration; therefore, the study of the mechanisms of Ca^2+^ accumulation in a muscle cell and its removal from it is important for understanding the processes of muscle contraction and relaxation.

Ca^2+^ enters the muscle cell from three sources: the extracellular space, the sarcoplasmic reticulum, and the mitochondria [1, 2, 7, 8]. In each of these sources, the concentration of ionized calcium is approximately 3-4 orders of magnitude higher than the Ca^2+^ concentration in the cytosol of the muscle cell in the basal state [1]. This concentration gradient is made possible by membranes: the plasma membrane (PM), which isolates the cell from the extracellular space and membranes covering the sarcoplasmic reticulum (SR) and mitochondria (MX). Membranes play a crucial role in calcium homeostasis because they contain protein structures that enable calcium cations to pass through the membrane in both directions. Ca^2+^ enters the cytosol via the PM, SR, and MX calcium channels, that allow calcium cations to pass down the concentration gradient (passive transport). Calcium pumps remove Ca^2+^ from the cytosol against a concentration gradient (active transport). Ion exchangers are also involved in the process of calcium removal from the cytosol [1, 3, 4, 7, 9].

The combination of passive and active transport mechanisms allows Ca^2+^ to redistribute quite easily between the cytosol, SR, MX, and the extracellular space, resulting in the establishment of a dynamic equilibrium in the cellular system. This enables the cell system to return to the initial (basal) state, which is characterized by strictly defined Ca^2+^ concentrations in cytosol, SR, MX and extracellular space after any stimulating action.

Researchers believe that there are several types of ion channels on the PM of smooth muscle cells, which enables Ca^2+^ to enter the cytoplasm. Most of them distinguish voltage-operated (VOC), which are also called voltage-depended or voltage-gated calcium channels, receptor-operated (ROC) and calcium channels that are sensitive to depletion of calcium stores in the SR (store-operated calcium channels or SOCs) [2, 4, 5, 7, 10-15]. Two types of calcium channels exist on the membrane of SR, each with corresponding receptors: ryanodine (RyR) and inositol 1,4,5-trisphosphate (IP_3_R) [5, 7, 10, 11, 16]. Both types of channels are Ca^2+^-dependent, and it is believed that Ca^2+^ channels are activated at low Ca^2+^ concentrations and inhibited at high concentrations [5, 7]. These channels are considered to differ in that RyR channels open in response to an increase in Ca^2+^ concentration in the cytosol caused by the opening of voltage-depended plasma membrane channels in response to plasma membrane depolarization [10, 17, 18], while IP_3_R channels are activated by inositol 1,4,5-trisphosphate, which is formed as a result of agonist action on receptor-operated channels of PM [7, 17].

Calcium pumps on the PM and on the membrane of SR are Ca^2+^-ATPases and are designated PMCA and SERCA, respectively [4, 8, 12, 16]. PMCA activity is strongly dependent on the cytosolic Ca^2+^ concentration (at low Ca^2+^ concentrations, about 100 nM), reaching a saturation level at Ca^2+^ concentrations of about 400 nM [8]. Another participant in Ca^2+^ removal from the cytosol is the Na^+^ / Ca^2+^ exchanger [16], but its productivity is lower than that of PMCA (it removes 25-30 % of all Ca^2+^ that enters the cytosol, versus 70-75 % removed by PMCA [4]).

In general, the process of changes of Ca^2+^ concentration in the intracellular space is currently presented as follows: activation of receptor-operated calcium channels leads to the appearance of inositol 1,4,5-trisphosphate in the cytosol, which opens the IP_3_R channels of SR, resulting in the rapid release of large amounts of Ca^2+^ from SR stores into the cytosol [17]; the activation of voltage-operated calcium channels, which occurs as a result of PM depolarization, leads to entry into the cytosol from the extracellular space of a certain amount of Ca^2+^, which, when interacting with the ryanodine receptors, opens RyR channels of SR, which also leads to rapid emptying of SR stores and, consequently, to a rapid increase in Ca^2+^ concentration in the cytosol. The latter process is called calcium-induced calcium release and abbreviated CICR [2, 9, 16, 17, 19]). The increase of Ca^2+^ concentration in cytosol activates both PMCA and SERCA, and Ca^2+^ starts to be pumped out of cytosol both into SR and extracellular space [17]. The depletion of the calcium stores of SR causes Ca^2+^ entry into the cell from the extracellular space through calcium channels of PM (the mechanism of this process is still unclear), which are called store-operated channels or SOCs [7, 15, 17, 20, 21]. This process is called store operated Ca^2+^ entry or SOCE [15, 16, 21] (which was earlier known as capacitive Ca^2+^ entry or CCE [7, 14, 22, 23]), and it allows to refill the SR stores with Ca^2+^ up to basal level. The current through the SOCs fixed using the patch-clamp method is called *I*_*CRAC*_ (Ca^2+^-release-activated Ca^2+^ current) [21, 24].

The latter processes reduce the Ca^2+^ concentration in the cytosol to the initial (basal) value, as well as refill the SR stores with Ca^2+^ to the basal level. Thus, the cellular system returns to its original state.

Of course, the picture considered is already considerably simplified. First of all, as mentioned earlier, mitochondria [1, 3, 4, 7, 9, 12, 16], which can rapidly release Ca^2+^ into the cytosol and rapidly pump it out from the cytosol, like SR, in this way take part in calcium homeostasis, as well as Na^+^ / Ca^2+^ exchanger of PM [1, 3, 4, 7, 9, 16], transporting Ca^2+^ from the cytosol into the extracellular space. There are also two other classes of ion channels on the PM of smooth muscle cell that are Ca^2+^-sensitive – K_Ca_ and Cl_Ca_ [7, 11], whose activation leads to depolarization of PM, as well as calcium-binding proteins in the cytosol and SR [4, 12]. However, even without this, the picture is quite complicated, since the detailed mechanism for almost any step has not been established, due to the fact that it is difficult to reproduce the physiological process under experimental conditions.

One of the most important issues is the process of calcium-induced calcium release from SR. Firstly, such attention is due to the importance of this process for medicine, particularly cardiology. It is now clear to researchers that an excessive increase in cytosolic Ca^2+^ concentration plays a major role in the development of myocardial damage during ischemic reperfusion and that Ca^2+^ release from SR is a critical process in the early phase of ischemia. Therefore, it is believed that any intervention in the process of emptying calcium stores in the SR in order to lower the rate of calcium release from the SR should be cardioprotective [9]. Secondly, this process is key to understanding the physiological role of Ca^2+^ in muscle contraction-relaxation.

Despite a large number of studies, much remains unclear in the mechanism of calcium-induced calcium release [7, 17]. It is still not entirely clear how Ca^2+^ opens and closes calcium channels of SR [5, 7], whether it only opens RyR channels or IP_3_R channels as well [4, 5, 7, 17]; how the avalanche-like Ca^2+^ release from SR is stopped [7, 9] and how the frequently observed periodic fluctuations of Ca^2+^ concentration in cytosol under agonist stimulation can be explained [9, 17].

Another equally important and interesting process is SOCE, i.e., the refilling of the SR store with calcium due to the opening of the SOCs. This process seems even more puzzling than CICR, since very synchronous interactions between structures (PM and SR) that are remote from each other and separated by the cytosol, often giving the impression that Ca^2+^ exchange between the extracellular space and SR occurs uninvolved cytosol. There is also much that is still unclear here: (*i*) what is the nature of SOCs [8, 14, 21, 25],(*ii*) how is the communication between SR and SOCs organized so that SOCs can sense when SR is emptied [2, 8], (*iii*) how SR stores are refilled with extracellular Ca^2+^, if in general refilling occurs when cytosolic Ca^2+^ concentration is near basal level and no longer changes substantially [8] and (*iv*) how SOCs sense when SR stores is completely filled with Ca^2+^ again to close [2].

Various assumptions have been made in scientific literature about possible mechanisms of activation and inhibition by calcium cations of channels and pumps on PM and SR [3, 5, 7, 8, 11, 12, 17, 24, 26, 27], about the mechanisms responsible for SOCE [2, 8, 12, 20-22, 24], theoretical models have been constructed to reproduce some of the observed experimental results [3, 6, 27-32], but so far the processes discussed above remain largely unexplained.

To better understand the CICR process, we developed and investigated a calcium-induced calcium release model.

Using modeling we have attempted, in particular, to find answers to the following questions:

- can cytosolic Ca^2+^ regulate the process of Ca^2+^ release from the SR?
- how the avalanche-like process of Ca^2+^ release from the SR, caused by an increase in the concentration of Ca^2+^ in the cytosol, can be stopped?
- whether it is necessary to close the calcium channels on PM with cytosolic Ca^2+^ to terminate the increase of Ca^2+^ concentration in the cytosol?
- what causes the appearance of oscillations of Ca^2+^ concentration in the cytosol under the agonist stimulation?
- how are SR stores refilled with Ca^2+^ from the extracellular space?
- can such refilling be carried out through the cytosol so that the Ca^2+^ concentration in the cytosol remains practically unchanged?
- can this process take place without SOCs?
- how can the complete filling of SR stores with Ca^2+^ have an effect on the closing of the SOCs?
- how can precisely coordinated processes occur in the cell, leading to the fact that after the termination of stimulation, the Ca^2+^ concentrations in the cytosol, SR and extracellular space return to the previous (basal) level (i.e., to the level that was before stimulation)?

## 2. Calcium-induced calcium release model

## 2.1 Model description

In order to find answers to these questions, we developed a mathematical model of the calcium-induced calcium release from SR, focusing primarily on smooth muscle cells. It was created on the basis of the model we studied earlier [33]. An essential addition to the former model is the presence of SR, which allowed us to study the Ca^2+^ concentration changes in cytosol, SR and extracellular space from the beginning of cell stimulation by a signal substance (agonist) up to complete return of the cell to the basal state when stimulation would be terminated.

Several similar models have been published in the literature [3, 6, 27-30, 32], the purpose of which was to achieve agreement with the experiment. Therefore, these models took into account practically all ideas about calcium homeostasis in the cell and covered all possible participants of this process. As a result, although there was some agreement between the model calculations and the results of real experiments, no answers to the above questions were found. The system was so complex that it was difficult to monitor the mutual influence of different processes and understand how they affect the process as a whole.

Therefore, we decided to make our model as simple as possible, but to take into account all necessary components, avoiding duplication and unnecessary complications. At the same time, the cell system should return to the previous state (which was before stimulation), as it happens in a living organism in norm. In this connection, our model considers a smooth muscle cell consisting of a plasma membrane isolating the cytosol from the extracellular space (with the cell volume equal to the volume of the extracellular space) with calcium channels (ROC) and a calcium pump located on it, as well as a sarcoplasmic reticulum with calcium channels (RyR) and calcium pump also located on its membrane. As a first approximation, we assumed that the concentration of calcium channels on PM and SR membrane is the same, just as the parameters of calcium pumps of PM and SR are the same. In the absence of stimulation, the cell is in a steady state (in the framework of the model it is an equilibrium state) characterized by a stable dynamic equilibrium with Ca^2+^ concentrations in the cytosol, SR and in the extracellular space equal to 100 nM, 1 mM and 1 mM, respectively (basal state).

Calcium channels on the outer side of the PM have receptors, whose interaction with a signaling substance that arrives in the extracellular space as a signal from other organs leads to the opening of the calcium channels of PM. We chose this way of stimulation (through action on receptors) because it is physiologically important and represents a way of therapeutic action on body organs and tissues (for example, on smooth muscles in their excessive contraction due to hypertension or asthma) [13].

There is information in the literature about experimental results showing that under conditions close to physiological ones, application of moderate concentrations of substances stimulating contraction leads to an increase in Ca^2+^ concentration in the cytosol of myocytes (the increase in Ca^2+^ concentration depends on the dose of physiologically active substances) and this increase is due to the extracellular cation pool [1]. It is also known that the conductance of voltage-depended channels can be regulated by neurotransmitters and hormones [1]. Nelson et al. note that vasoconstrictors such as NE and serotonin can depolarize the arterial membrane, causing Ca^2+^ entry into the cell through voltage-depended channels, or can increase Ca^2+^ entry into the smooth muscle cell through PM with no significant depolarization [34]. Based on these data, the authors consider the possibility that NE receptor-dependent channels and voltage-depended calcium channels of PM are actually one and the same. The article [14] also discusses such a possibility.

When modeling the interaction of calcium channel receptors located on the PM with a signal substance, we based on the fact that in the body, in response to stimulation, a substance that is a signal-transmitter substance (a signal substance), such as a hormone, is secreted or synthesized with which special enzymes interact immediately, and as a result this signal substance is quickly enough completely decomposed [35]. Such enzyme catalyzed transformation of the signaling substance is incorporated in our model, but the signaling substance in the model is injected immediately at its maximum concentration. The disappearance of the signaling substance under the action of the enzyme can also, in the first approximation, be considered as the injection of the pharmacological drug (which is not subject to enzymatic transformation) and its further removal from the zone of influence on the cell by blood flow.

Calcium channels on the cytosolic side of the SR also have receptors that can interact with cytosolic Ca^2+^, thus opening these channels for Ca^2+^ release from the SR into the cytosol.

In our model, the calcium channels on the SR have ryanodine receptors (RyR). The characteristics of the calcium channels were specially chosen by us because they allow us to investigate a complex process with the involvement of a minimum required set of components. Therefore, in our model, stimulation is carried out through receptor-operated channels (because this is the most physiological way of stimulation), with Ca^2+^ (the main regulator of muscle contraction) entering the cell and initiating the calcium-induced calcium release from SR.

To simplify the model, it does not include mitochondria, but only SR, because MX somewhat duplicates SR by releasing Ca^2+^ into the cytosol and pumping it out of the cytosol. The mitochondrial Ca^2+^ uniporter is known to have a lower affinity for Ca^2+^ than SERCA and is probably only important when cytosolic Ca^2+^ is elevated above 0.5 μM [36]. From this point of view, SR can be considered, within the framework of the model, as an “integral” structure (SR+MX), which contains the store of intracellular Ca^2+^, which can be released into the cytosol and pumped back out again. Of course, such an approximation may to some extent affect the shape of the kinetic dependences, but, in our opinion, it will not be able to distort the qualitative picture of the process.

Our model considers SR not as a single structure but as a set of 100 autonomous structures (compartments, vesicles, cisternae) of the same size, each with calcium channels and a calcium pump on its membrane – this approach is based on the literature data on SR in smooth muscle [2, 5]. Note that in the model the SR can also be represented as a single structure (the corresponding model parameter must be changed for this purpose).

Buffer proteins were also not considered in the model because their presence (in physiological concentrations) should not qualitatively change the kinetic dependences, and there is also evidence in the literature that these proteins have little effect both on the value of the maximum calcium concentration in the cytosol and on the time to reach this maximum [29]. We also did not take into account Na^+^ / Ca^2+^ exchanger because the model does not consider Na^+^ homeostasis in the cell. The same applies to K^+^, so the model does not consider changes in PM potential.

Thus, the smooth muscle cell in which the process of calcium-induced calcium release takes place can be represented as follows:

1. a smooth muscle cell has a volume *V*_*cell*_ and is surrounded by a plasma membrane with an area *S*_*PM*_ on which there is a constant (equal to *S*_*PM*_.[*R*]_*all*_) number of calcium channels (where [*R*]_*all*_ is the surface concentration of calcium channels of the plasma membrane, i.e., the number of calcium channels per unit area of myocyte membrane);
2. there are a certain number (*n*_*r*_) of SR cisternae in the cell, the volume and surface area of each of them are equal to *V*_*r*_ and *S*_*r*_, respectively. On the surface of each SR cisterna there is a constant (equal to *S*_*r*_.[*RR*]_*all*_) number of calcium channels (where [*RR*]_*all*_ is the surface concentration of calcium channels of the SR, i.e., the number of calcium channels per unit surface area of the membrane of SR cisterna);
3. in the steady state there is a small Ca^2+^ flux into the cell through the calcium channels of PM (which we will call the slow basal Ca^2+^ flux through the closed calcium channels of PM (SBF-PM) not to be confused with the passive Ca^2+^ entry into the cytosol through the open Ca^2+^ channels of PM) [1, 34], which is compensated by its pumping out from the cytosol into the extracellular space by the calcium pump of PM. Similarly, in the steady state there is a small Ca^2+^ flux into the cytosol from the SR (slow basal Ca^2+^ flux through closed Ca^2+^ channels of the SR (SBF-SR) or leak [37]). Ca^2+^ passes into cytosol from the SR also through calcium channels opened by Ca^2+^ present in the cytosol (at 100 nM). The total flow of Ca^2+^ from the SR into the cytosol is compensated by its pumping out of the cytosol into the SR by calcium pumps of SR cisternae (each SR cisterna has its own calcium pump);
4. an average Ca^2+^ concentration is rapidly established in the cytosol, so all SR cisternae equally participate in maintaining calcium homeostasis in the myocyte (the percentage of open channels and their throughput capacity are the same for all cisternae; the calcium pumps of each cisterna are also characterized by the same parameters of Michaelis constants and the limiting rate), so at any time the Ca^2+^ concentration inside SR cisternae is the same;
5. signal substance *A* binds to calcium channel receptor *R* on the outer surface of the PM resulting in opening of the channels and increasing their throughput capacity;
6. there are *RR* receptors on the surface of SR cisternae [19] that can bind cytosolic Ca^2+^ [3], resulting in opening of the channels and significantly increasing their throughput capacity. Open calcium channels of SR can have a significantly higher throughput capacity than open calcium channels of PM;
7. signal substance *A* is destructed by the enzyme, resulting in a gradual decrease in its concentration, which leads to a decrease in the amount of signal substance bound to the receptors, and, consequently, a decrease in the number of open calcium channels of the PM;
8. open calcium channels of PM can also be closed due to blockage by cytosolic Ca^2+^ (“Ca^2+^-dependent inactivation”) [1, 2, 38].

### 2.2 Description of the steady state (basal state)

Since there are indications in the literature that Ca^2+^ can itself reversibly inactivate various calcium channels of PM [1, 2, 5, 38], the model took into account that calcium channels of PM have Ca^2+^ binding sites on the cytoplasmic side. When calcium channels on the PM open, Ca^2+^ enters the cell. Subsequently, due to the increased concentration of Ca^2+^ in the cytosol, Ca^2+^ binds with these sites, leading the calcium channels to close. We believe that cytosolic Ca^2+^ can interact with such calcium channel sites whether the channels are open or not. In the steady state (in the absence of signaling substance), calcium channels are in the closed state *R*, passing Ca^2+^ at a slow rate (SBF-PM [1, 34]). At the same time, we consider that cytosolic Ca^2+^ (with a basal concentration of 100 nM) are able to bind to the calcium channel binding sites on the inner side of the PM, forming the *RCa* complex (the channels remain closed) and their surface concentration in steady state is [*RCa*]_*b*_:

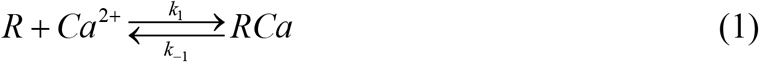

Regardless of the state of the channel (*R* or *RCa*), it has a fixed (slight) throughput capacity (SBF-PM) [1, 34], which is characterized by the *D*_*PM*_ coefficient. The passive calcium flux into the cytoplasm from the extracellular space in the basal state (*J*_*p,PM,b*_) is determined by Fick’s law:

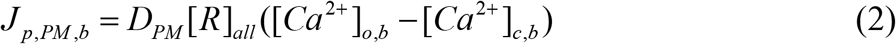

where [*Ca*^*2+*^]_*o,b*_ and [*Ca*^*2+*^]_*c,b*_ are concentrations of Ca^2+^ in the extracellular space and in the cytosol in the basal state, respectively.

In the steady state (basal state), the active Ca^2+^ flux through the membrane into the extracellular space from the cytosol is exactly the same in magnitude but reversed in direction. It is generated by the calcium pump of PM (*J*_*a,PM,b*_), and is described by the Michaelis-Menten equation within the framework of this model:

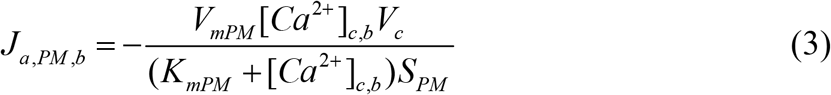

where *V*_*mPM*_ is the limiting rate of the calcium pump of PM, *K*_*mPM*_ is the Michaelis constant relative to Ca^2+^, *V*_*c*_ is the volume of smooth muscle cell excluding SR volume (i.e. cytosol volume).

Due to the equality of Ca^2+^ fluxes through the PM inside the smooth muscle cell and out of the myocyte, we obtain the first equilibrium equation:

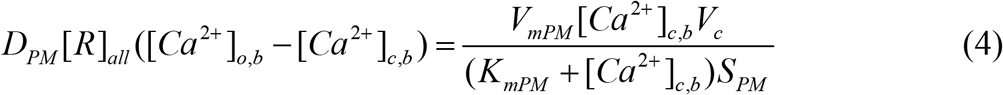

Since under steady state conditions the cytosol contains Ca^2+^ at a concentration of 100 nM, some of these cations will bind to calcium channel receptors on SR (*RR*) [3]

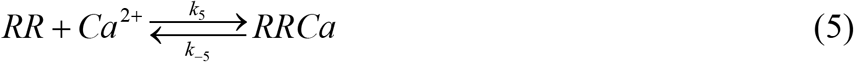

which will inevitably lead to the fact that part of the calcium channels of SR will be open, allowing Ca^2+^ to pass from SR to cytosol [37, 39]. The passive Ca^2+^ flux from SR to cytosol for the steady state (as the sum of the SBF-SR and the Ca^2+^ flux through the calcium channels of SR which are open in the basal state) can also be written using Fick’s law:

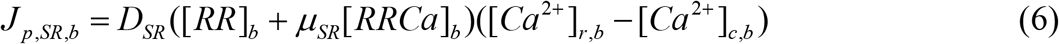

where [*RR*]_*b*_, [*RRCa*]_*b*_ are surface concentrations of calcium channels on the SR membrane in the closed and open states, respectively; [*Ca*^*2+*^]_*r,b*_ are basal Ca^2+^ concentrations in the SR, *μ*_*SR*_ is the conductance gain coefficient of the calcium channel of SR.

The active Ca^2+^ flux through the SR membrane from the cytosol to the SR of the same magnitude, but in the opposite direction, is generated by the calcium pump of SR, and can also be described by the Michaelis-Menten equation:

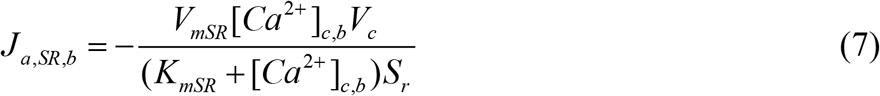

where *V*_*mSR*_ is the limiting rate of the calcium pump of SR, *K*_*mSR*_ is the Michaelis constant of the calcium pump of SR with respect to Ca^2+^.

Thus, using the condition of equality of Ca^2+^ fluxes through the SR membrane, we obtain the second equilibrium equation:

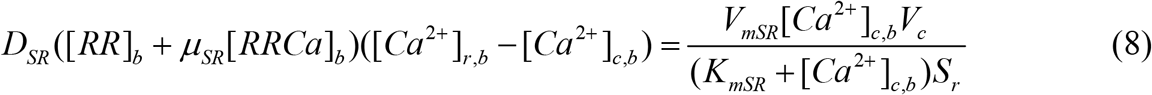

To determine the number of open and closed calcium channels on the PM and SR membrane in the basal state, we use the equations derived from the equilibrium states of reactions (1) and (5):

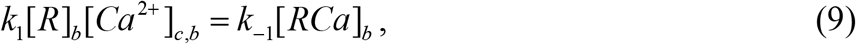

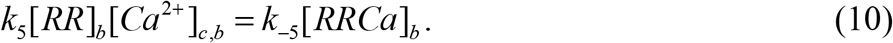

Considering equalities [*R*]_*b*_ + [*RCa*]_*b*_ = [*R*]_*all*_, [*RR*]_*b*_ + [*RRCa*]_*b*_ = [*RR*]_*all*_, we can derive from equations (9), (10) equations for the surface concentrations of calcium channels on PM and on SR membrane in the basal state

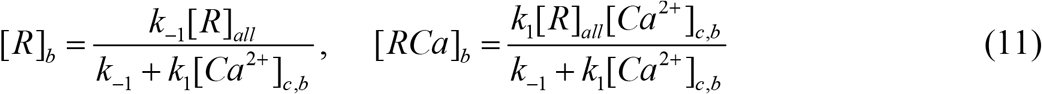

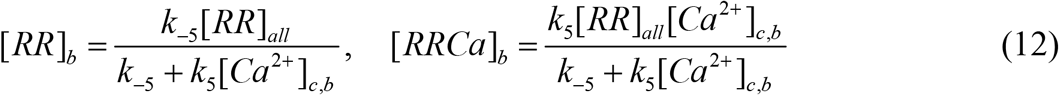

Thus, equations (4), (8), (11), (12) describe the steady state (basal state) of the smooth muscle cell system.

### 2.3 Description of non-stationary processes

The non-stationary process begins when the signaling substance (agonist) *A* appears in the extracellular space (the volume of which per myocyte is taken as *V*_*o*_) and interacts with calcium channel receptors located on the outer side of PM [40]:

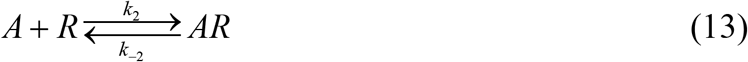

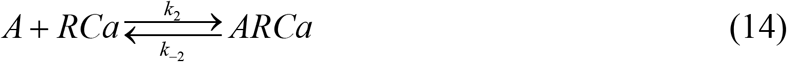

Let us assume that the rate constants of reactions do not depend on the state of the channel (*R* or *RCa*). Signaling substance *A*, once it appears in the extracellular space, immediately degraded by the enzyme *F*. First, it binds with enzyme *F*, producing an enzyme-substrate complex *AF*

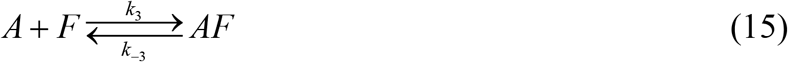

which, then, decomposes with the regeneration of enzyme *F* and the formation of the reaction product*P*

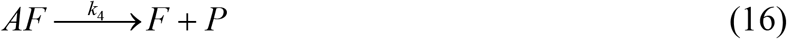

This produces a gradual disappearance of signaling substance *A* outside the cell.

Thus, calcium channels are present on the PM in four states: open *AR*; closed *R*; closed inactivated *RCa*; open inactivated *ARCa*. We assume that the calcium channels of PM in the *R, RCa*, or *ARCa* states all have the same throughput capacity with coefficient *D*_*PM*_ (i.e., they are in fact closed channels), whereas the throughput capacity of the calcium channel of PM in the *AR* state (i.e., open channel) is significantly higher (*μ*_*PM*_ .*D*_*PM*_). So, the agonist amplifies the throughput capacity of calcium channels of PM not inactivated by Ca^2+^ by *μ*_*PM*_ times.

Similarly, we assume that the calcium channels of SR in the closed state (*RR*) have a throughput capacity with coefficient *D*_*SR*_, whereas the throughput capacity of the open calcium channel of SR (*RRCa*) is significantly higher (*μ*_*SR*_.*D*_*SR*_). So, activation of the calcium channels of SR by cytosolic Ca^2+^ increases the throughput capacity of these channels by *μ*_*SR*_ times.

Reactions (1), (5), (13-16) represent the chemical aspect of the process. Based on these reactions, we can calculate concentrations of open and closed channels on PM and SR, as well as concentrations of *A, F, FA*, and *P*. Let’s write down the corresponding kinetic equations:

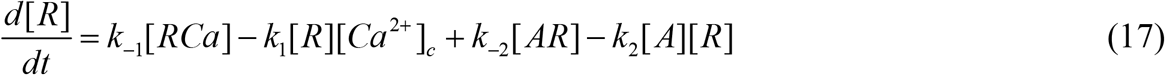

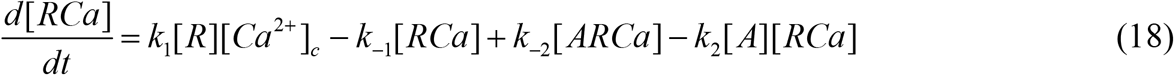

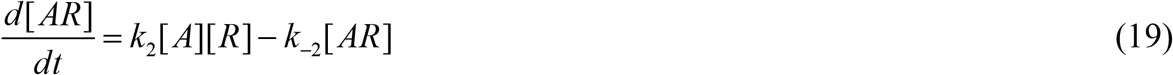

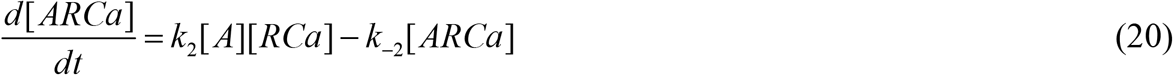

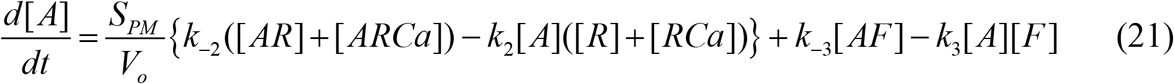

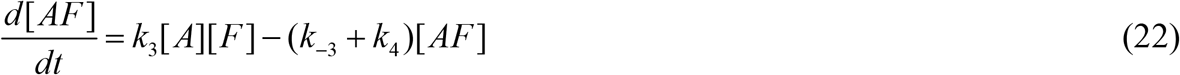

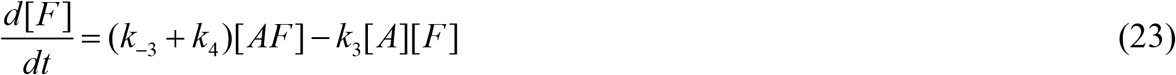

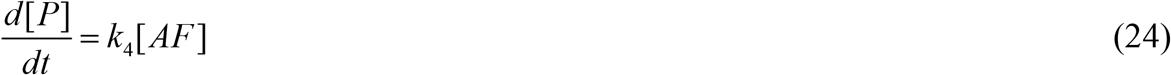

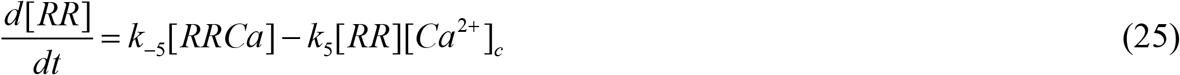

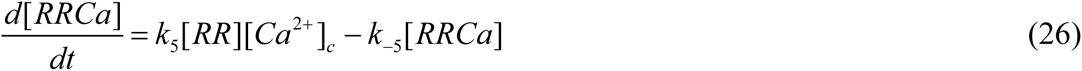

Now we need to calculate Ca^2+^ concentrations in the extracellular space, in the cytosol, and in the SR. To do this, let us first write the equations for Ca^2+^ fluxes through PM and SR membrane. The passive Ca^2+^ flux into the cytosol from the extracellular space (*J*_*p,PM*_) is proportional to the throughput capacity of calcium channels, the number of open channels and the gradient of Ca^2+^ concentrations on both sides of PM:

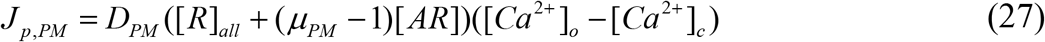

The active Ca^2+^ flux from the cytosol to the extracellular space (*J*_*a,PM*_) is determined by the calcium pump of PM:

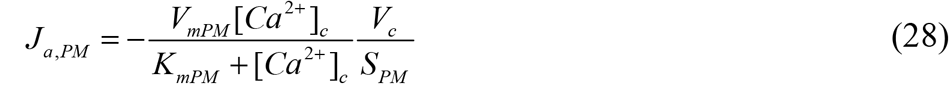

The passive Ca^2+^ flux into the cytosol from SR (*J*_*p,SR*_) is proportional to the throughput capacity of calcium channels, the number of open channels, and the gradient of Ca^2+^ concentrations on both sides of the SR membrane:

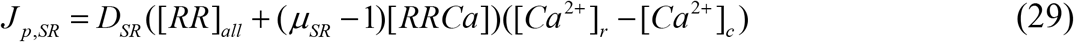

The active Ca^2+^ flux from the cytosol to the SR (*J*_*a,SR*_) is determined by the calcium pump of SR:

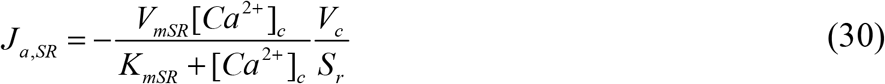

Note that equations (27) and (29) (similar equations are often reported in the literature to describe passive transport, for example [1, 3, 6, 27-31]) indicate that calcium channels, both PM and SR, are permeable to Ca^2+^ in both directions. While under physiological conditions it is unlikely that the Ca^2+^ concentration in the cytosol would be higher than in the SR or in the extracellular space, under experimental conditions such cases are often observed. As an example, Ca^2+^ is completely removed from the extracellular space or SR stores are completely emptied. Under such conditions, active and passive transport can be carried out in one direction, and this must be taken into account in such studies.

As a consequence of these fluxes, Ca^2+^ is distributed between the cytosol, the SR, and the extracellular space. Therefore, knowing the flux values, we can calculate how much Ca^2+^ is in the extracellular space, cytosol, and SR at any time point. The change of Ca^2+^ mass in the extracellular space *m*_*o*_*(t)* and in the SR *m*_*r*_*(t)* will be described by the following equations:

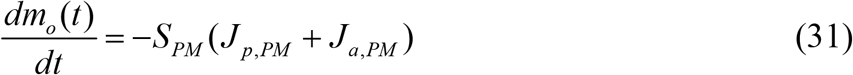

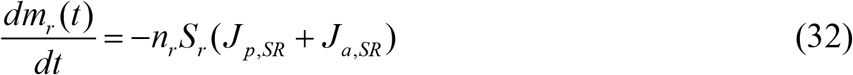

To calculate the change of Ca^2+^ mass in the cytosol *m*_*c*_*(t)*, along with the fluxes, the binding of cytosolic Ca^2+^ to receptors on the PM and the SR membrane due to reactions (1) and (5) must also be considered:

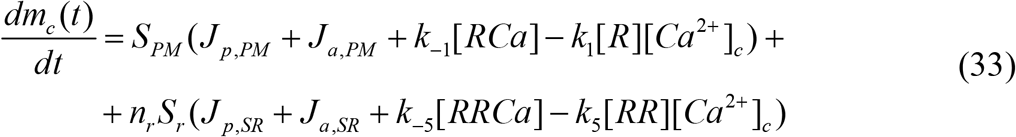

Based on equations (32), (27), and (28), let us write a differential equation to calculate the extracellular Ca^2+^ concentration:

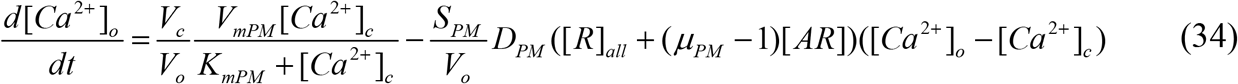

From equations (31), (29) and (30) we derived a differential equation to calculate the Ca^2+^ concentration in SR:

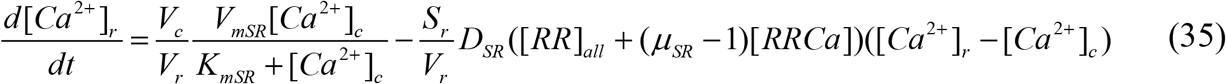

Finally, from equations (33) and (27-30) we obtained a differential equation to calculate the cytosolic Ca^2+^ concentration:

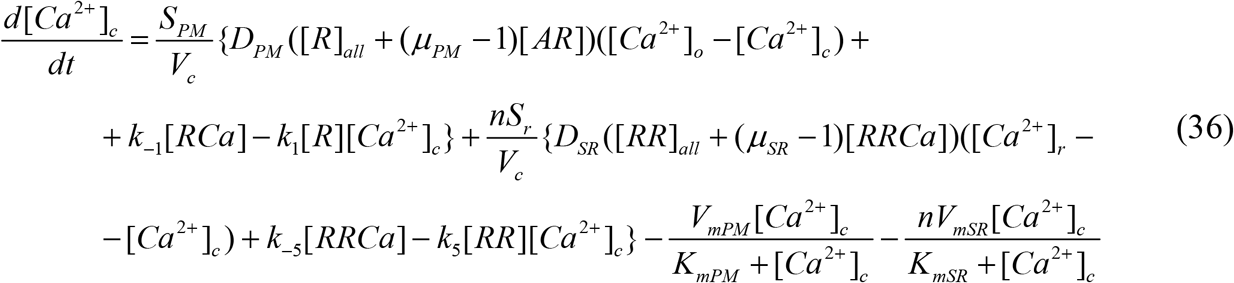

Thus, the mathematical model of non-stationary processes of Ca^2+^ transport through PM and membrane of SR induced by signal substance *A* is a Cauchy problem for the system of ordinary differential equations (17-26) and (34-36). The initial conditions are determined by the initial concentrations of substances: [*R*] | _*t* =0_ = [*R*]_*b*_, [*RCa*] | _*t* =0_ = [*RCa*]_*b*_, [*AR*] |_*t* =0_ = 0, [*ARCa*] | *t* =0 = 0, [*A*] | _*t* =0_ = [*A*]_0_, [*AF*] |_*t* =0_ = 0, [*F*] | _*t* =0_ = [*F*]_0_, [*P*] |_*t* =0_ = 0, [*RR*] | _*t* =0_ = [*RR*]_*b*_, [*Ca*^2+^]_*o*_ |_*t* =0_= [*Ca*^2+^]_*o,b*_ ?, [*Ca*^2+^]_*r*_ |_*t* =0_= [*Ca*^2+^]_*r,b*_, [*Ca*^2+^]_*c*_?|_*t* =0_ = [*Ca*^2+^]_*c,b*_, [*RRCa*] |_*t* =0_ = [*RRCa*]_*b*_, where the surface concentrations [*R*]_*b*_, [*RCa*]_*b*_, and [*RR*]_*b*_, [*RRCa*]_*b*_ in the basal state are determined by formulas (11) and (12), respectively.

Note that the parameters *D*_*PM*_, [*R*]_*all*_, [*Ca*^2+^]_*c,b*_, [*Ca*^2+^]_*o,b*_, *V*_*mPM*_, *K*_*mPM*_ are not arbitrary values, but are related by equation (4). Therefore, by setting the values of [*R*]_*all*_, [*Ca*^2+^],_*c,b*_ [*Ca*^2+^]_*o,b*_, *V*_*mPM*_, *K*_*mPM*_, we unambiguously determine the throughput capacity coefficient *D*_*PM*_ of calcium channel of PM (and the rate of SBF-PM). Similarly, relation (8) allows us to calculate the throughput capacity coefficient *D*_*SR*_ of calcium channels of SR in the basal state. Thus, the basal state in our developed model makes it possible to “tune” the cellular system using throughput capacity coefficients of calcium channels so that passive and active transport through PM and the SR membrane are equilibrated, allowing basal concentrations to be maintained in cytosol, SR and extracellular space (100 nM, 1 mM and 1 mM, respectively). This basal state adjustment plays a very important role, since any stimulation of the cellular system generates a driving force to return the system to the basal state.

The mathematical models of the steady state and non-stationary state of the myocyte are interrelated, because after the ending of the signal substance action, the cell system again returns to the same steady state (basal state) as before the beginning of the stimulation. In the steady state (basal state), the cell can remain for any length of time without visible changes in macroscopic parameters, in particular, Ca^2+^ concentration in cytosol and SR while Ca^2+^ concentration in the extracellular space remains unchanged (although the calcium pumps on PM and SR membrane are continuously functioning, pumping out Ca^2+^, which permanently permeates into cytosol from extracellular space and SR).

The fact that the cell system can return to the same state as it was before the stimulation is an essential factor for describing the kinetics of cyclic processes in smooth muscle cells and understanding the driving forces of these processes. In addition, the proposed model illustrates the mechanism of initiation of transmembrane Ca^2+^ transport using signal substance *A*, which can be degraded by the enzyme. This approach makes it possible to naturally initiate the process of Ca^2+^ transport through PM and, at the same time, to limit the duration of action of the signaling substance on this process.

Since the steady state is characterized by strictly defined values of Ca^2+^ concentrations in the cytosol, SR and in the extracellular space, and the cell system always restores the values of these concentrations when returning to the basal state, it is more suitable to compare the current concentrations with the basal level: *C*_*cyt*_ = [*Ca*^*2+*^]_*c*_ */* [*Ca*^*2+*^]_*c,b*_ – how many times the Ca^2+^ concentration in cytosol changed in comparison with the basal level; *C*_*ret*_ = [*Ca*^*2+*^]_*r*_ */* [*Ca*^*2+*^]_*r,b*_ – how many times the Ca^2+^ concentration in SR changed in comparison with the basal level; *C*_*out*_ = [*Ca*^*2+*^]_*o*_ */* [*Ca*^*2+*^]_*o,b*_ – how many times the Ca^2+^ concentration in extracellular space changed in comparison with the basal level.

For the extracellular space, where the calcium concentration changes insignificantly during the stimulation process, the difference between the current and basal Ca^2+^ concentrations was also used: *D*_*out*_ = [*Ca*^*2+*^]_*o*_ *–* [*Ca*^*2+*^]_*o,b*_. Relative values were also used for signal substance concentration *C*_*A*_ = [*A*] / [*A*]_0_, for determination of the number of calcium channels opened with signal substance on PM (in relation to the total number of calcium channels on PM) *R*_*open*_ = [*AR*] / [*R*]_*all*_, and for determination of the number of open calcium channels of SR (in relation to the total number of calcium channels of SR) *RR*_*open*_ = [*RRCa*] / [*RR*]_*all*_.

To test the hypothesis that cytosolic Ca^2+^ at high concentrations can interact with open channels on the cytosolic side of PM, closing them and terminating Ca^2+^ entry into the cytosol from the extracellular space, we examined how many PM calcium channels are closed by cytosolic Ca^2+^ of those that were opened by the signaling substance:

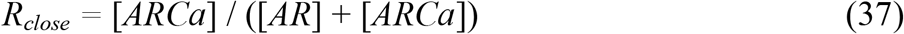

In addition to the Ca^2+^ fluxes that were determined earlier (equations (27-30)), attention was also paid to the total Ca^2+^ fluxes through the PM (38) and the SR membrane (39)

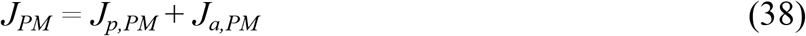

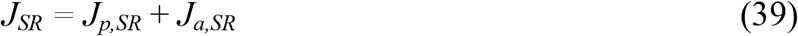

Each of these fluxes are expressed in mol.dm^-2.^s^-1^. However, since these fluxes refer to a unit area of the corresponding membrane, it is difficult to estimate the contribution of each of them in the process of Ca^2+^ accumulation in the cytosol. Therefore, let us calculate all fluxes for the entire surface area of the corresponding membrane and obtain the absolute values of fluxes through the entire membrane (PM or SR), expressed in moles of Ca^2+^ transported per second. Afterwards the total Ca^2+^ flux through PM into cytosol from extracellular space will be expressed by equation (40), the total Ca^2+^ flux through PM into extracellular space from cytosol by equation (41), the total Ca^2+^ flux through SR membrane into cytosol from SR by equation (42), and the total Ca^2+^ flux through SR membrane from cytosol into SR by equation (43)

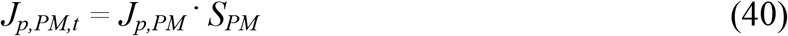

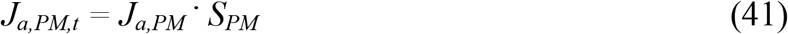

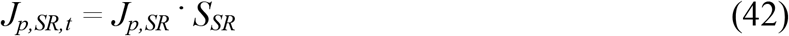

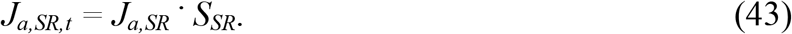

Such fluxes can already be compared with each other, and it can be written down that the flux into the cytosol by passive Ca^2+^ transport (from the extracellular space and SR) is equal:

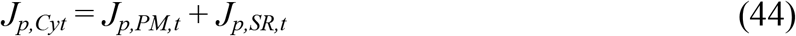

The flow from the cytosol (to the SR and extracellular space) due to the functioning of calcium pumps, in turn, is expressed by the formula:

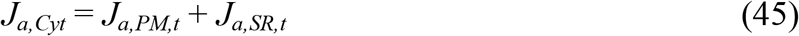

The sum of fluxes *J*_*p,cyt*_ and *J*_*a,cyt*_ will show the magnitude and direction of the total Ca^2+^ flux relative to the cytosol, i.e., the balance of Ca^2+^ entering and leaving the cytosol:

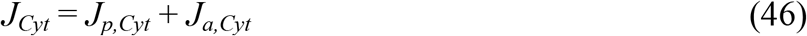

A positive *J*_*Cyt*_ value means that Ca^2+^ enters the cytosol, while a negative value means that Ca^2+^ is pumped out of the cytosol.

### 2.4 Comparison of the model parameter values with literature data

Let us first evaluate to what extent the process parameters within our model (presented in Table 1) correspond to the experimental data obtained in the study of real smooth muscle preparations. The average volume of smooth muscle is usually *V*_*sm*_ = 2000-4000 μm^3^ [1, 29], the sarcolemma area is *S*_*sm*_ = 1200-6000 μm^2^ [1], and the cell geometry is such that the *S*_*sm*_ / *V*_*sm*_ ratio is in range of 1 to 3 [1]. In our model, these parameters are 4000 μm^3^, 8000 μm^2^, and 2, respectively.

**Table .** Initial values of the kinetic model parameters.

| Symbol | Parameter meaning | Value | Dimension |
| --- | --- | --- | --- |
| $[Ca^{2+}]_{o,b}$ | $Ca^{2+}$ concentration in the extracellular space | 1 | mM |
| $[Ca^{2+}]_{r,b}$ | $Ca^{2+}$ concentration in the sarcoplasmic reticulum | 1 | mM |
| $[Ca^{2+}]_{c,b}$ | $Ca^{2+}$ concentration in the cytosol | 100 | nM |
| $[R]_{all}$ | surface concentration of calcium channels (receptors) on PM | 0.01 | nmole/dm<sup>2</sup> |
| $[RR]_{all}$ | surface concentration of calcium channels (receptors) on the SR membrane | 0.01 | nmole/dm<sup>2</sup> |
| $[A]_0$ | concentration of the signal substance introduced into the extracellular space | 0.01–10 | μM |
| $[F]_0$ | concentration of the enzyme that degrades the signal substance | 20–30 | nM |
| $V_o$ | volume of extracellular space per myocyte | 4000 | μm<sup>3</sup> |
| $V_{cell}$ | intracellular space volume (including SR volume) | 4000 | μm<sup>3</sup> |
| $V_c$ | cytosolic volume | 3996 | μm<sup>3</sup> |
| $n_r$ | number of SR cisternae in the myocyte | 100 | pieces |
| $V_r$ | the volume of one SR cisterna | 0.04 | μm<sup>3</sup> |
| $S_r$ | the surface area of the membrane of one SR cisterna | 0.5656 | μm<sup>2</sup> |
| $V_{SR}$ | total volume of all SR cisternae | 4 | μm<sup>3</sup> |
| $S_{SR}$ | total surface area of the membranes of all SR cisternae | 56.56 | μm<sup>2</sup> |
| $S_{PM}$ | surface area of plasma membrane of myocyte | 8000 | μm<sup>2</sup> |
| $\mu_{PM}$ | conductance gain coefficient of calcium channel of PM | 10 | |
| $\mu_{SR}$ | conductance gain coefficient of calcium channel of SR | 50–2000 | |
| $V_{mPM}$ | limiting rate of the calcium pump of PM | 200 | nM/s |
| $V_{mSR}$ | limiting rate of the calcium pump of SR | 200 | nM/s |
| $K_{mPM}$ | Michaelis constant of the calcium pump of PM | 100 | nM |
| $K_{mSR}$ | Michaelis constant of the calcium pump of SR | 100 | nM |
| $k_1$ | rate constant of $Ca^{2+}$ binding on the cytosolic side of calcium channels of PM ($Ca^{2+}$-dependent inactivation) | $1 \cdot 10^9$ | 1/M's |
| $k_{-1}$ | rate constant of unblocking of calcium channels on the cytosolic side of PM, due to $Ca^{2+}$ uncoupling | $3 \cdot 10^4$ | 1/s |
| $k_2$ | rate constant of signaling substance binding by receptors of calcium channel on PM (receptor-dependent activation of calcium channels of PM) | $5 \cdot 10^8$ | 1/M's |
| $k_{-2}$ | rate constant of inactivation of calcium channels of PM due to termination of the action of the signal substance | $3 \cdot 10^4$ | 1/s |
| $k_3$ | rate constant of binding of a signal substance by an enzyme that destroys it, with the formation of an enzyme-substrate complex | $2 \cdot 10^8$ | 1/M's |
| $k_{-3}$ | the rate constant of decay of the enzyme-substrate complex with signal substance regeneration | $3 \cdot 10^3$ | 1/s |
| $k_4$ | rate constant of decomposition of the enzyme-substrate complex to products | $1 \cdot 10^2$ | 1/s |
| $k_5$ | rate constant of $Ca^{2+}$ binding to receptors of SR membrane ($Ca^{2+}$-dependent activation of SR channels) | $1 \cdot 10^8$ | 1/M's |
| $k_{-5}$ | rate constant of inactivation of calcium channels SR, due to the detachment of calcium from the receptors of calcium channels of SR | $1 \cdot 10^4$ | 1/s |

Smooth muscle cells usually occupy 65 % of the muscle preparation [1]. In our model, the cell volume is equal to the volume of the extracellular space, so that the above ratio is 50 %. SR volume according to the literature is a few percent of smooth muscle cell volume (2-5 % [1], 6 % [11], 1.5-7.5 % [4, 41]) and surface area is 10-20 % of smooth muscle cell surface area [1]. In our model, the ratio of volumes and areas is an order of magnitude lower – 0.1 % and 0.7 %, respectively.

The concentration of Ca^2+^ in the cytosol is usually approximately 10^-7^ M [6, 8, 16, 26, 27, 29, 31, 35, 38, 41-43], as in our model. In the extracellular space, the Ca^2+^ concentration is usually 1-2 mM [1, 2, 4-6, 8, 16, 34, 35, 38, 43]: in our model [*Ca*^*2+*^]_*r,b*_ = 10^-3^ M. The following Ca^2+^ concentrations in SR were found in the literature: 0.1-0.3 mM [8, 31, 35, 41, 44], 0.3-1.0 mM [45], and 1.0-2.0 mM [27, 43], and in our model [*Ca*^*2+*^]_*r,b*_ = 10^-3^ M.

According to the data, myocytes contain on average from 10^6^ (for the cardiac cell) [42] to 1.5.10^6^ [3] ryanodine receptors. Assuming that the SR area is between 120 and 600 μm^2^ [1], the concentration of RyR on the SR membrane should be about 1600-12500 μm^-2^. Up to 10^6^ [24] CRAC channels can be expressed on the PM of a single lymphocyte cell. It can be assumed that along with receptor-operated calcium channels, 1.5.10^6^ calcium channels are present on the PM of the smooth muscle cell. Assuming that the myocyte plasma membrane area is between 1200 and 6000 μm^2^ [1], the concentration of calcium channels of all types on the PM should be approximately 250-1250 μm^-2^. In our model, the concentration of receptors on both the SR membrane and the PM is about 600 μm^-2^. The total number of calcium channels on the PM is 4816000 and on the SR membrane is 34050.

It is noted in the existing literature that part of the cytosolic Ca^2+^ is localized on the inner surface of PM [1, 2, 38]. Experimental measurements of the dissociation constant give a Ca^2+^ affinity value with PM (particularly for myometrium) of 5.6-8.7.10^-5^ M [1]. In [3] the following values of the kinetic constants were used for the calculations: the rate constant for the formation of the Ca^2+^ complex with PM is 1.10^8^ M^-1.^s^-1^ and the rate constant for the dissociation of this complex is 1300 s^-1^, which gives a dissociation constant of 1.3.10^-5^ M; in [38] these values were respectively 2.2.10^6^ M^-1.^s^-1^, 0.6 s^-1^ and 2.7.10^-7^ M. The corresponding values for our model are 1.10^9^ M^-1.^s^-1^, 3.10^4^ s^-1^, and 3.10^-5^ M. Usually, the binding of agonists to receptors is characterized by dissociation constants approximately 10^-7^-10^-8^ M [40]. In our model, the corresponding constant is 6.10^-5^ M. To calculate the kinetics of Ca^2+^ binding to the SR, the following constants were used in [3]: the forward reaction rate constant is 1.10^8^ M^-1.^s^-1^ and the reverse one is 60 s^-1^, which gives a dissociation constant of 6.10^-7^ M. In our model, the corresponding constants are 1.10^8^ M^-1.^s^-1^, 1.10^4^ s^-1^, and 1.10^-4^ M.

It has been found in experimental work that for the calcium pump of myometrial sarcolemma *K*_*m*_ = 300-500 nM and *V*_*m*_ = 4-5 μM.min^-1^ or 60-80 nM.s^-1^ (with a Hill coefficient that can vary from 1 to 2) [1]. Shannon et al. [3] used values of *K*_*m*_ = 500 nM and *V*_*m*_ = 2.2 μM.s^-1^ (with a Hill coefficient of 1.6), and in our model *K*_*m*_ = 100 nM and *V*_*m*_ = 200 nM.s^-1^ (with a Hill coefficient of 1.0). For the calcium pump of SR *K*_*m*_ = 0.3-1.2 μM, and the Hill coefficient can also take different values – in particular, its values of 1.2 and 1.7 were found for the SR fraction isolated from the smooth muscle cell of the pig coronary artery [1]. In [3], *K*_*m*_ = 246 nM and *V*_*m*_ = 286 μM.s^-1^ (with a Hill coefficient of 1.787) were used for calculations, and in our model (the same as for the calcium pump of PM), *K*_*m*_ = 100 nM and *V*_*m*_ = 200 nM.s^-1^ (with a Hill coefficient of 1.0). Here we emphasize that in our model the calcium pump is present in every SR cisterna, i.e., there are 100 pieces in total.

An important generalizing parameter is the value of the slow Ca^2+^ flux into the cell from extracellular space in the basal state (i.e., SBF-PM) – this flux for smooth muscle cells is usually from 1.10^-15^ to 1.10^-14^ mol Ca^2+^/cm^2.^s [1]. In our model with the parameter values given in Table 1, in the basal state the flux into the cell through the PM (as well as the flux from the cell through the PM) is *J*_*p,PM,b*_ = – *J*_*a,PM,b*_ = 4.995.10^-15^ mol Ca^2+^/cm^2.^s, which corresponds to *J*_*p,PM,t,b*_ = – *J*_*a,PM,t,b*_ = 3.996.10^-19^ mol Ca^2+^/s i.e. 240560 calcium ions pass through PM in one and the other direction in one second (the same number of calcium ions are in the cytosol in the steady state, so that they can be completely renewed in one second; as can be seen, this flux is quite powerful, although it is called slow). Therefore, the total Ca^2+^ flux through PM in the basal state *J*_*PM,b*_ = *J*_*p,PM,b*_ + *J*_*a,PM,b*_ = 0. In the steady state, the basal Ca^2+^ concentration in the cytosol (100 nM) is the reason that part of this Ca^2+^ binds to SR receptors, opening calcium channels of SR (9.99.10^-15^ mol/dm^2^ or 0.0999 % of all calcium channels on the SR membrane). These open channels, along with the SBF-SR, provide a passive Ca^2+^ flux into the cytosol from the SR *J*_*p,SR,b*_ = 7.0648.10^-11^ mol Ca^2+^/cm^2.^s (which corresponds to *J*_*p,SR,t,b*_ = 3.996.10^-17^ mol Ca^2+^/s or 24055920 calcium ions per second – a total of 2408000 calcium ions in the sarcoplasmic reticulum in the basal state), which is compensated by an equal in magnitude Ca^2+^ flux from the cytosol to SR, due to the functioning of the calcium pump of SR. It is known from the literature that the calcium flux through the SR membrane of skeletal muscle in the basal state is 1.10^-13^ -1.10^-12^ mol Ca^2+^/cm^2.^s [1].

In the basal state, a part of Ca^2+^ from the cytosol is also consumed for binding to calcium channels on the inner side of the PM. Only 0.33 % of all calcium channels on the PM are bound to calcium cations on the inner side of the PM in the basal state (at the parameters given in Table 1).

### 2.5 Slow basal flow from extracellular space

There are different points of view in the existing literature regarding the existence of SBF-PM. While some authors register such flux in the studied cells [1, 4, 29, 34], others do not find it [46], arguing that Ca^2+^ ions have an extremely low permeability through the PM. However, in our opinion, it is unlikely from the thermodynamic point of view that with such a huge gradient of Ca^2+^ concentrations (almost four orders of magnitude) there would be no diffusive transfer of Ca^2+^ across the PM and the SR membrane. Apparently, even the presence of a charge on the PM cannot prevent this, since not calcium in the form of cations but molecules or ionic pairs that have no charge can penetrate through the cell membrane [47].

This flux can be seen in an increase in Ca^2+^ concentration in the cytosol when Ca^2+^ concentration in the extracellular space increases [46, 48, 49]. The presented model was developed for such myocytes in which SBF-PM exists. SBF-PM gives the model new fundamental properties: being a basic flux on which other fluxes are superimposed under stimulation conditions, it determines the parameters of functioning of calcium pump of PM, together with it is responsible for maintaining the basal level of Ca^2+^ concentration in cytosol and allows the system of calcium homeostasis to function fully without positive and negative feedbacks regulating functioning of calcium pump of PM.

The kinetics of Ca^2+^ concentration changes in the cytosol is determined by four fluxes [29, 30]: (*i*) total passive Ca^2+^ flux into the cytosol from the extracellular space through agonist-opened calcium channels of PM combined with SBF-PM (*J*_*p,PM,t*_); (*ii*) total passive Ca^2+^ flux into the cytosol from SR, including the calcium-induced calcium release (*J*_*p,SR,t*_); (*iii*) total active Ca^2+^ flux from the cytosol to the SR due to the functioning of the calcium pump of SR (*J*_*a,SR,t*_); (*iv*) total active Ca^2+^ flux from the cytosol to the extracellular space, the value of which is determined by the parameters of the calcium pump of PM (*J*_*a,PM,t*_).

With the chosen parameters of calcium pumps of PM and SR, their initial rates in the basal state are 0.5.*V*_*m*_, allowing for both upward and downward modulation.

## 3 Study of a model of calcium-induced calcium release in a smooth muscle cell

First of all, we should note that, of course, it is quite difficult to fully investigate such a complex nonlinear model. As can be seen from Table 1, the model includes about thirty variables, the values of most of which may vary over a wide range, depending on the studied smooth muscle tissue, and for some (such as reaction rate constants) the experimental values in general are unknown. However, as a result of studies, two characteristic features of the model behavior were found, which can be briefly called a single cytosolic Ca^2+^ transient mode [37, 50] and an oscillatory mode [37, 51]. Any ratios of the variables always lead to one or the other mode (at least, we have not discovered any other modes).

### 3.1 Single cytosolic Ca^2+^ transient mode: the buffering role of SR

When the signal substance is in the extracellular space, it binds to the receptors of the PM, thus opening the calcium channels of the PM. Ca^2+^ from the extracellular space then enter the cytosol and involves SR in the process of Ca^2+^ accumulation in the cytosol.

Depending on the ratio of the model parameters, SR can act both as a passive participant in the process of Ca^2+^ accumulation in the cytosol of the smooth muscle cell and play a major role in this process. If the conductance gain coefficient of calcium channel of SR is slightly higher than conductance gain coefficient of calcium channel of PM (50 versus 10), the process of Ca^2+^ accumulation in cytosol occurs practically thanks to extracellular Ca^2+^ (Fig. 1, curve 1), and SR plays the role of buffer accumulating part of Ca^2+^ that entered from cytosol (Fig. 1, curve 7). In this case, the Ca^2+^ concentration in SR does not decrease below the basal level during the whole process of stimulation (while the concentration of extracellular Ca^2+^ does not increase above the basal level during the whole process of stimulation (Fig. 1, curve 4); that is, the Ca^2+^ concentration in cytosol and in SR is changed exclusively on account of extracellular Ca^2+^).

**Fig. 1.**
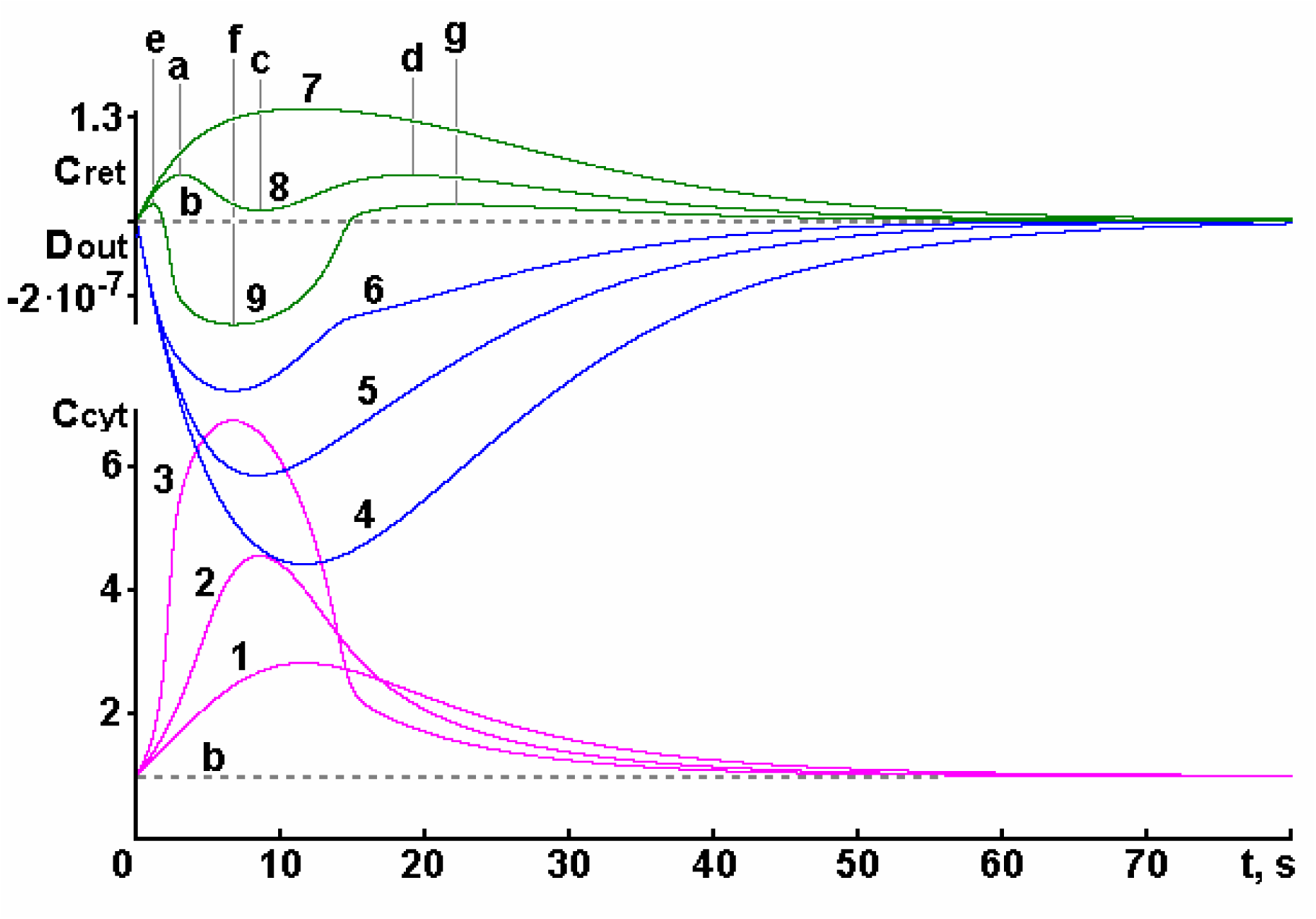
Dependences of Ca^2+^ concentration in cytosol *C*_*cyt*_ = [*Ca*^*2+*^]_*c*_ */* [*Ca*^*2+*^]_*c,b*_ (curves 1, 2, 3), in extracellular space *D*_*out*_ = [*Ca*^*2+*^]_*o*_ *–* [*Ca*^*2+*^]_*o,b*_ (curves 4, 5, 6) and in SR *C*_*ret*_ = [*Ca*^*2+*^]_*r*_ */* [*Ca*^*2+*^]_*r,b*_ (curves 7, 8, 9) on time *t* at *μ*_*SR*_ = 50 (curves 1, 4, 7), *μ*_*SR*_ = 200 (curves 2, 5, 8), *μ*_*SR*_ = 400 (curves 3, 6, 9). [*A*]_0_ = 10 μM, [*F*]_0_ = 20 HM. The values of other parameters are given in Table 1. The corresponding basal levels are indicated by the letter ***b***.

At the same time, the Ca^2+^ concentration in SR first increases (up to 11.652 s, Fig. 1, curve 7; at this stage part of the Ca^2+^ from the cytosol is pumped into SR, slowing the increase in the cytosolic Ca^2+^ concentration) and then slowly decreases to the basal level in the process of establishing a steady state in the cell system (at this step part of the Ca^2+^ previously pumped into SR is relocated into the cytosol, slowing the decrease in the cytosolic Ca^2+^ concentration). At the time point when the cellular system returns to the basal state, all Ca^2+^ that was pumped into SR in the first step leaves SR and Ca^2+^ concentration in it returns to the basal level. The same takes place in the cytosol and extracellular space.

Looking at the distribution of Ca^2+^ entering the cell from the extracellular space between cytosol and SR, it was found that the first portions of Ca^2+^ almost completely remain in cytosol, then the proportion of Ca^2+^ accumulated in cytosol decreases, reaching a minimum at 0.864 s (22.7 % of the total Ca^2+^ that enters from the extracellular space), and then increases again, reaching 35 % at 11.51 s (0.06 s earlier than the maximum Ca^2+^ concentration in the cytosol is observed, see Fig. 1, curve 1). In the further, the percentage of Ca^2+^ accumulated in the cytosol decreases again.

As can be seen in Fig. 1 (curve 1) the maximum cytosolic Ca^2+^ concentration is observed at 11.569 s (*t*_*max*_ = 11.569 s) and it is 2.83 times higher than the basal level (*C*_*cyt,max*_ = 2.83). The Ca^2+^ flux into cytosol from extracellular space *J*_*p,PM*_ reaches the maximal level (2.25 times more than basal level) at 0.0002 s when the number of calcium channels of PM opened by signal substance is maximal (*R*_*open*_ = 0.139). In the further it gradually decreases, but the active Ca^2+^ transport through the PM *J*_*a,PM*_ keeps increasing until it reaches its maximum value at *t*_*max*_ = 11.569 c. At this time the maximum Ca^2+^ content in the cytosol is observed, which also causes the closure of the maximum number of calcium channels open by the signaling substance on the PM (*R*_*close*_ *=* 0.0093), the opening of the maximum number of calcium channels on the SR membrane (*RR*_*open*_ = 0.0028) and the maximum rate of Ca^2+^ pumping into SR (*J*_*a,SR*_ by 48 % exceeds basal level).

Both passive and active Ca^2+^ transport through the PM (fluxes *J*_*p,PM*_ and *J*_*a,PM*_) decrease after time *t*_*max*_ = 11.569 s, but they decrease at different rates. Equality of the fluxes *J*_*p,PM*_ and *J*_*a,PM*_ (and, respectively, *J*_*PM*_ = 0) is observed at 11.595 s (which is 0.026 s later than *t*_*max*_). At the same time the minimum Ca^2+^ concentration in the extracellular space is also reached (*C*_*out*_ = 0.999443, see Fig. 1, curve 4). The maximum rate of Ca^2+^ pumping out of the cytosol into the extracellular space (*J*_*PM*_) is achieved only at 22.53-22.54 s, when *J*_*a,PM*_ is greater than *J*_*p,PM*_ by about 20 %. The highest rate of Ca^2+^ entry into the cytosol from the extracellular space (*J*_*PM*_) is observed at 0.0002 s, when *J*_*p,PM*_ is approximately 2.25 times greater than *J*_*a,PM*_.

Interestingly, the maximum Ca^2+^ flux from SR to cytosol (*J*_*p,SR*_) is observed not at the time of opening the maximum number of calcium channels of SR, as one would expect, but somewhat later, at 11.609 s. This is because in addition to the number of open channels, the difference in Ca^2+^ concentrations in SR and cytosol also affects the flux value. The maximum Ca^2+^ concentration in SR (Fig. 1, curve 7) is observed at 11.652 s (*C*_*ret*_ = 1.36). At this point, *J*_*p,SR*_ = *J*_*a,SR*_ and *J*_*SR*_ = 0. The maximum rate of Ca^2+^ pumping out of the cytosol into the SR (*J*_*SR*_) is reached at 0.107 s, and the highest rate of Ca^2+^ flux from the SR into the cytosol (*J*_*SR*_) is reached at approximately 25.9 s.

If the fluxes due to passive (*J*_*p,Cyt*_) and active transport (*J*_*a,Cyt*_) are examined separately, we can see that both fluxes increase after stimulation, but *J*_*p,Cyt*_ increases faster, reaching a maximum of 11.481 s (*J*_*p,Cyt*_ = 5.96425.10^-17^ mol/s), whereas *J*_*a,Cyt*_ becomes maximum at *t*_*max*_ = 11.569 s (*J*_*a,Cyt*_ = 5.96420.10^-17^ mol/s). As a result, the total Ca^2+^ flux into the cytosol *J*_*Cyt*_ initially increases rapidly, reaching a maximum (*J*_*Cyt*_ = 4.9662.10^-19^ mol/s) at 0.00014 s then it begins to decrease (*J*_*Cyt*_ = 0 at *t*_*max*_ = 11.569 s), and after *t*_*max*_ the Ca^2+^ pumping out of the cytosol commences to prevail, reaching its maximum at 19.146 s (*J*_*Cyt*_ = –3.75.10^-20^ mol/s).

Therefore, from the time when the signaling substance begins to work and until the steady state is established (agonist stimulation process), one cycle of filling/emptying of SR occurs: initially, part of the Ca^2+^ from the cytosol is pumped into the SR (pumping continues until the Ca^2+^ concentration in the cytosol reaches its maximum value), thus slowing the rapid increase in cytosolic Ca^2+^ concentration, and then, as Ca^2+^ begins to be pumped out of the cytosol, the SR slowly begins to return stored Ca^2+^ into the cytosol, slowing the rate of decrease in the cytosolic Ca^2+^ concentration.

We can conclude that when Ca^2+^ enters the cytosol from the extracellular space slowly enough, the calcium pumping a part of Ca^2+^ into the SR, succeed in reducing the Ca^2+^ concentration in the cytosol to a level that is insufficient to provoke a massive release of Ca^2+^ from SR.

### 3.2 Single cytosolic Ca^2+^ transient mode: the role of SR in enhancing the calcium signal

When the conductance gain coefficient of the calcium channel of SR is increased up to 200 (Fig. 1, curves 2, 5, 8), we can see that during the agonist stimulation process the SR performs two cycles of filling/emptying of SR. Moreover, the role of SR in the process of Ca^2+^ accumulation in cytosol of the smooth muscle cell changes dramatically: it is no longer limited to the role of a buffer but starts to actively participate in the formation of the calcium signal in the cytosol. In the first step (up to label ***a*** in Fig. 1) SR (as in the previous case) pumps the part of Ca^2+^ that entered the cytosol from the extracellular space: the maximum rate of Ca^2+^ pumping into SR (*J*_*SR*_) is reached at 0.12 s, resulting in a minimum rate of Ca^2+^ entry into the cytosol (*J*_*Cyt*_ = 1.24623.10^-19^ mol/s), whereas the maximum rate of Ca^2+^ entry into the cytosol is observed at 0.00014 s (*J*_*Cyt*_ = 4.9689.10^-19^ mol/s). However, even before reaching the maximum Ca^2+^ concentration in the cytosol, Ca^2+^ begins to leave the the SR, moving into the cytosol. The influence of SR becomes prominent (i.e., *J*_*SR*_ becomes positive) at 3.138 s (label ***a*** in Fig. 1), when the Ca^2+^ concentration in the SR reaches a maximum *C*_*ret*_ = 1.1473 (Fig. 1, curve 8). Maximum rate of Ca^2+^ flux from SR to cytosol (*J*_*SR*_) is reached at 5.138 s and as a result the rate of Ca^2+^ flux into cytosol increases, reaching a maximum at 4.41 s (*J*_*Cyt*_ = 2.7984.10^-19^ mol/s), causing a sharp increase in cytosolic Ca^2+^ (although total passive Ca^2+^ flux into cytosol *J*_*p,Cyt*_ continues to increase up to 8.329 s, when it reaches its maximum). This leads to a change in the shape of the left branch of the kinetic dependence of the cytosolic Ca^2+^ concentration: it becomes S-shaped (Fig. 1, curve 2).

After 4.41 s, the total Ca^2+^ flux into the cytosol (*J*_*Cyt*_) diminishes and becomes negative at 8.497 s, i.e. *J*_*p,Cyt*_ becomes less than *J*_*a,Cyt*_ (at this time *J*_*a,Cyt*_ becomes maximal and the Ca^2+^ concentration in the cytosol reaches its maximum *C*_*cyt,max*_ = 4.57 (Fig. 1, curve 2) at a moment earlier, at *t*_*max*_ = 8.496 c). At 8.56 s, the Ca^2+^ concentration in SR becomes minimal (but higher than the basal level – *C*_*ret*_ = 1.033), and this moment (label ***c*** in Fig. 1, curve 8) can be considered as the beginning of the second cycle of filling/emptying of SR by Ca^2+^. During this step, Ca^2+^ again begins to be pumped out of the cytosol into SR at an increasingly accelerating rate until at 12.39 s the *J*_*SR*_ reaches its maximum rate and at 12.803 s the total Ca^2+^ flux from the cytosol reaches its maximum (*J*_*Cyt*_ = –1.370.10^-19^ mol/s). Later, the rate of Ca^2+^ pumping into SR gradually decreases until at 18.762 s the *J*_*SR*_ becomes positive (at the same time the maximum amount of Ca^2+^ accumulates in the SR again (*C*_*ret*_ = 1.1474, see Fig. 1, curve 8). From this point (label ***d*** in Fig. 1) the final step begins: when the Ca^2+^ concentration in the cytosol is relatively low (*C*_*cyt*_ = 2.24), the Ca^2+^ stored in the SR begins transporting into the cytosol, slowing the decrease in the cytosolic Ca^2+^ concentration. The maximum rate of Ca^2+^ flux into the cytosol from the SR (*J*_*SR*_) is observed at 27.3 s, after which it slowly decreases. Thus, the first cycle of filling/emptying of SR lasts from the beginning of stimulation until 8.56 s (label ***c*** in Fig. 1), when Ca^2+^ is first pumped out of the cytosol into SR (up to label ***a*** in Fig. 1) and then it returns to the cytosol from SR (from label ***a*** to label ***c*** in Fig. 1). The second cycle begins at 8.56 s (label ***c*** in Fig. 1) and continues until the system returns to the basal state: during this time, Ca^2+^ is first pumped out of the cytosol into the SR again (from label ***c*** to label ***d*** in Fig. 1) and then it returns to the cytosol (from label ***d*** to the end of the process in Fig. 1).

We can see from Fig. 1 (curve 8) that, as in the previous case (curve 7), during the agonist stimulation process, the Ca^2+^ concentration in SR does not decrease below the basal level, i.e., SR contributes to the increase in the cytosolic Ca^2+^ concentration actually at the expense of Ca^2+^ pumped into SR during the previous step of the agonist stimulation process, rather than from its initial (basal) calcium reserves. However, unlike the first case (with *μ*_*SR*_ = 50), when SR works in antiphase with the plasma membrane during the whole agonist stimulation process (decreasing the cytosolic Ca^2+^ concentration when PM works to increase this concentration, and, conversely, increasing the Ca^2+^ concentration in the cytosol, when Ca^2+^ is pumped out of the cytosol by the calcium pump of PM), in the second case (with *μ*_*SR*_ = 200) SR begins to work synchronously with PM during a certain time (in the middle of the agonist stimulation process), amplifying its action (their joint activity allows Ca^2+^ to accumulate in the cytosol from 3.138 to 8.39 s and then leave the cytosol between 8.56 and 18.76 seconds).

If the distribution of Ca^2+^ entering the cell from the extracellular space between cytosol and SR is considered, then in this case initially Ca^2+^ completely remains in cytosol, later the percentage of Ca^2+^ accumulated in cytosol is decreasing (because some Ca^2+^ is pumped into SR), reaching a minimum at 0.58 s (30.5 % of the total Ca^2+^ entered from the extracellular space), and again begins to increase, reaching 92 % at 8.55 s (this increase from 30.5 to 92 % is achieved by the Ca^2+^ pre-stored in SR). Subsequently, the percentage of Ca^2+^ accumulated in the cytosol decreases again.

However, it is known from the literature that because of Ca^2+^ release into the cytosol, the SR is emptied. This phenomenon is observed only at higher values of the conductance gain coefficient of calcium channel of SR. At the value of this coefficient equal to 400, the Ca^2+^ release from SR becomes so significant that the Ca^2+^ concentration in SR decreases markedly below the basal level (the maximum decrease *C*_*ret*_ = 0.663 is observed at 6.7 s, see Fig. 1, curve 9): an avalanche-like release of Ca^2+^ from SR begins. In this case, the maximum Ca^2+^ concentration in the cytosol is *C*_*cyt,max*_ = 6.76 at *t*_*max*_ = 6.663 s (Fig. 1, curve 3). Analysis of Fig. 1 allows to conclude that with the increase of the conductance gain coefficient of the calcium channels of SR the maximum amount of accumulated Ca^2+^ in SR and the pumping time (up to reaching the maximum Ca^2+^ concentration in SR) decreases in the first cycle of filling/emptying of SR (*C*_*ret*_ = 1.1473 at 3.138 s for *μ*_*SR*_ = 200 (point ***a*** on curve 8) and *C*_*ret*_ = 1.0509 at 1.121 s for *μ*_*SR*_ = 400 (point ***e*** on curve 9)), and pumping in the second cycle occurs later and later (*C*_*ret*_ = 1.1474 at 18.762 s for *μ*_*SR*_ = 200 (point ***d*** on curve 8) and *C*_*ret*_ = 1.0513 at 21.888 s for *μ*_*SR*_ = 400 (point ***g*** on curve 9)). The results for *μ*_*SR*_ = 50 (*C*_*ret*_ = 1.36 and *t* = 11.652 s) can be interpreted as a condition where the first and second cycles have not yet separated.

### 3.3 Appearance of cytosolic Ca^2+^ concentration oscillations

It can be seen from Fig. 2 that with a further increase in the conductance gain coefficient of the calcium channel of SR a qualitative change in the kinetics of the process is observed – the cell system switches to oscillatory mode.

**Fig. 2.**
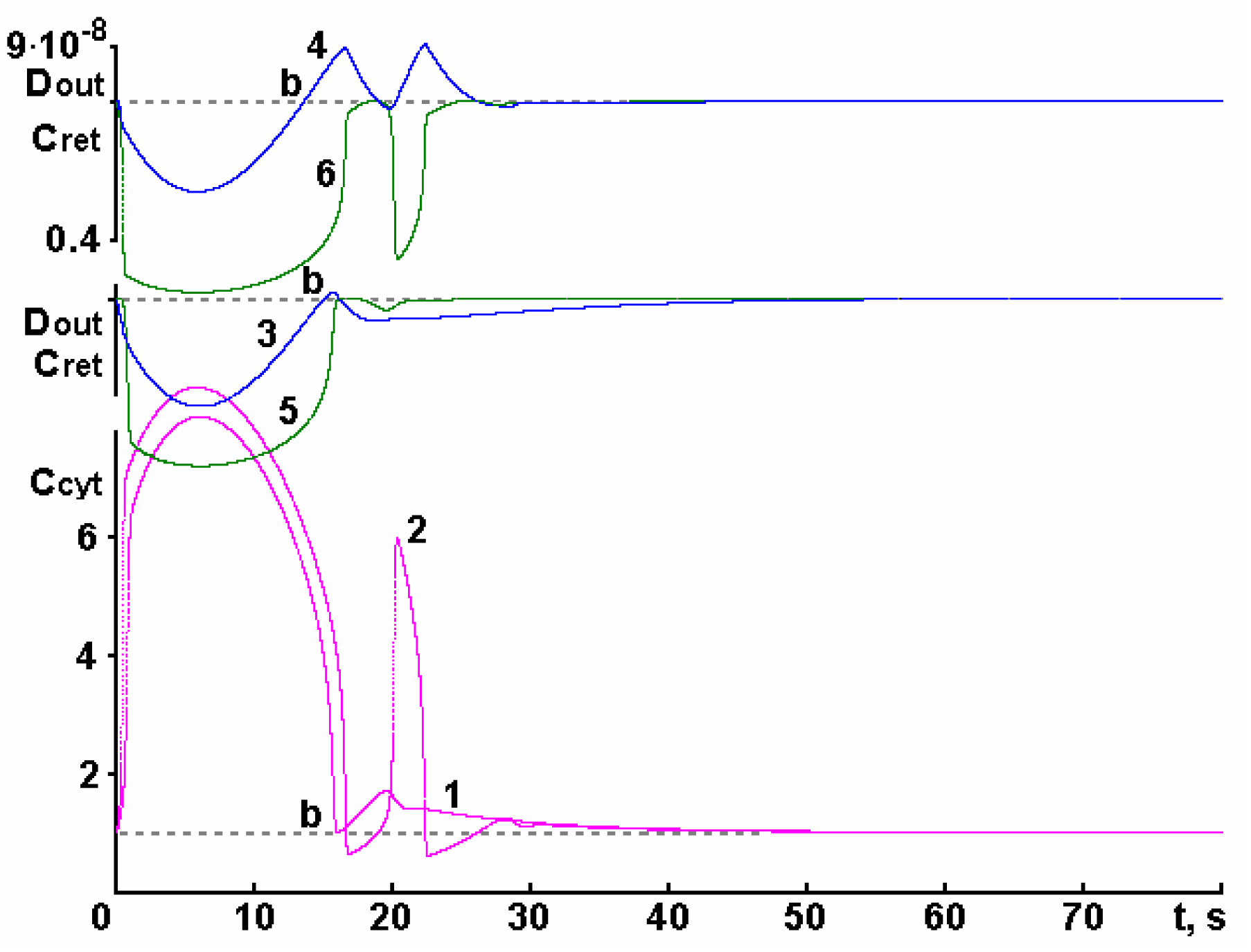
Dependences of Ca^2+^ concentration in cytosol *C*_*cyt*_ = [*Ca*^*2+*^]_*c*_ */* [*Ca*^*2+*^]_*c,b*_ (curves 1, 2), in the extracellular space *D*_*out*_ = [*Ca*^*2+*^]_*o*_ *–* [*Ca*^*2+*^]_*o,b*_ (curves 3, 4) and in SR *C*_*ret*_ = [*Ca*^*2+*^]_*r*_ */* [*Ca*^*2+*^]_*r,b*_ (curves 5, 6) on time *t* at *μ*_*SR*_ = 800 (curves 1, 3, 5), *μ*_*SR*_ = 1200 (curves 2, 4, 6). [*A*]_0_ = 10 μM, [*F*]_0_ = 20 HM. The values of other parameters are given in the Table 1. The corresponding basal levels are marked by the letter ***b***.

Moreover, the number of oscillations increases with the increase of the conductance gain coefficient of the calcium channels of SR. The process of appearance of new oscillations has the following scheme: at first, the small rapidly damped oscillations appear (curves 1, 3, 5), and then, with the increase of the conductance gain coefficient of the calcium channels of SR, initially the amplitude of the first oscillation increases to a certain level (curves 2, 4, 6), then the second, and so on (however, it is very difficult to explain why the kinks appear in certain places of the smooth kinetic dependences, which then turn into peaks – we can only state that it happens due to the nonlinearity of the system).

A series of large-amplitude oscillations is followed by several rapidly decaying oscillations with much smaller amplitudes. There are cases when during the whole the process, the Ca^2+^ concentration in SR and in the extracellular space practically does not exceed the corresponding basal level (Fig. 2, curves 5 and 3, respectively), but more typically the process proceeds so that the Ca^2+^ concentration both in the SR and in the extracellular space varies within a relatively wide range, both exceeding the basal level and decreasing below it (similar to curve 4 in Fig. 2).

### 3.4 Periodic oscillation mode

It is known from the literature that at low (physiological) concentrations of the active substance (in particular, hormones) the cytosolic calcium signal appears as repeated discrete peaks (spikes) [8]. Fig. 3 shows such a process. Let us note at once that the dependence of cytosolic Ca^2+^ concentration on time (curve 1) shown in Fig. 3 corresponds well in its shape, amplitude, and frequency to the experimentally obtained dependencies given in the articles [11, 52, 53].

**Fig. 3.**
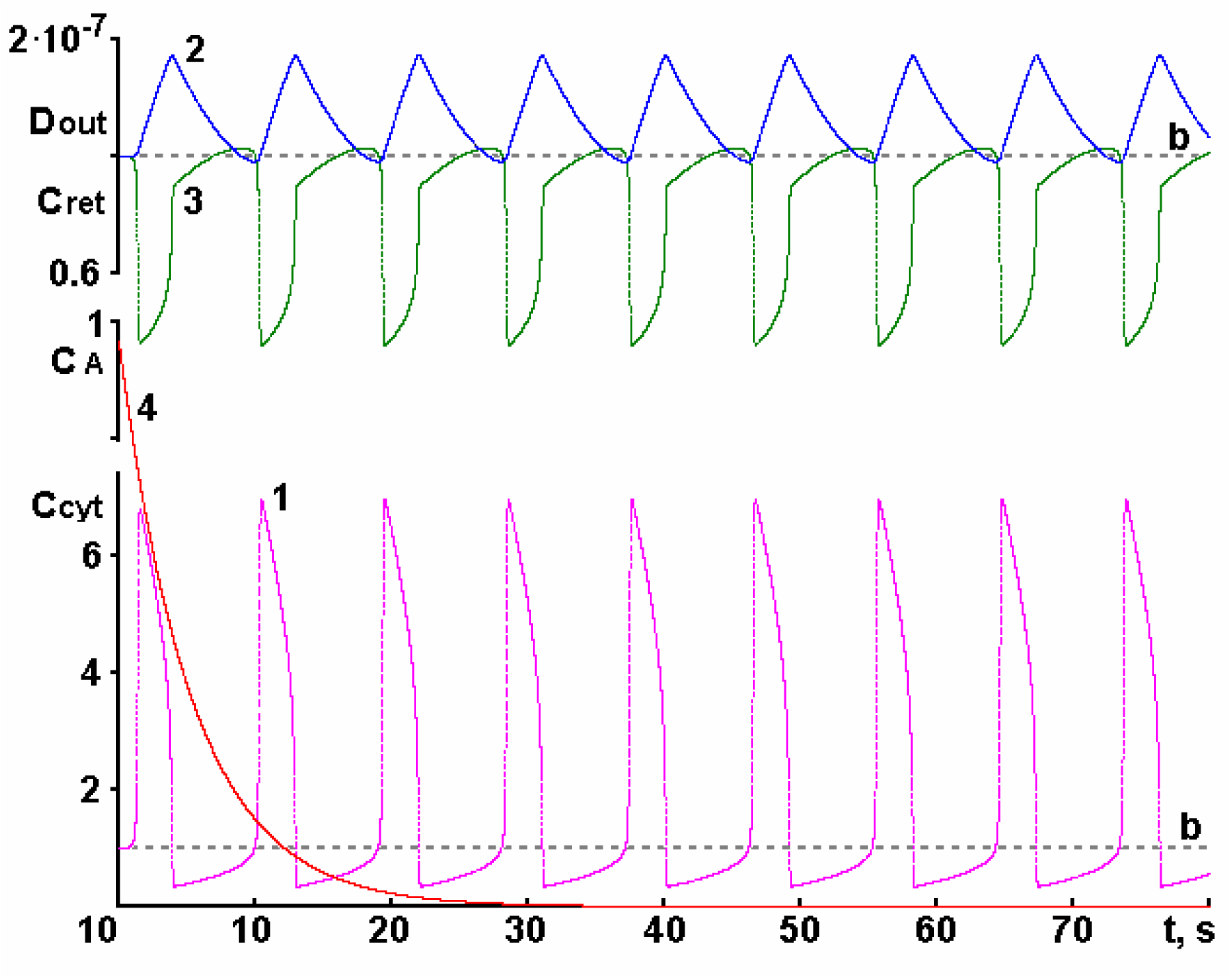
Dependences of Ca^2+^ concentration in cytosol *C*_*cyt*_ = [*Ca*^*2+*^]_*c*_ */* [*Ca*^*2+*^]_*c,b*_ (curve 1), in extracellular space *D*_*out*_ = [*Ca*^*2+*^]_*o*_ *–* [*Ca*^*2+*^]_*o,b*_ (curve 2), in SR *C*_*ret*_ = [*Ca*^*2+*^]_*r*_ */* [*Ca*^*2+*^]_*r,b*_ (curve 3) on time *t* and the change in signal substance concentration *C*_*A*_ = [*A*] / [*A*]_0_ (curve 4) over time at *μ*_*SR*_ = 2000, [*A*]_0_ = 10 HM, [*F*]_0_ = 30 HM. The values of other parameters are given in the Table 1. The corresponding basal levels are marked with the letter ***b***.

The authors of the article [52] investigating the rhythmic increases of cytosolic Ca^2+^ distinguish three phases in the obtained curves. In phase 1, there is a slow gradual increase in the cytosolic Ca^2+^ concentration up to a certain critical level (approximately 20 % of the amplitude of the cytosolic calcium signal), after which a rapid increase in the cytosolic Ca^2+^ concentration begins. According to the authors, this phase reflects some kind of rhythm-driven process. In phase 2, the cytosolic Ca^2+^ concentration increases rapidly until it reaches its peak value. As suggested by the authors, this is a consequence of Ca^2+^ release from the SR. In phase 3, the cytosolic Ca^2+^ concentration decreases from the maximum level to practically the basal level. The authors believe that this phase is a consequence of Ca^2+^ pumping out of the cytosol by the calcium pump of SR. They note that for rabbit portal vein the frequency of oscillations was 0.02-0.1 Hz, i.e., 1 oscillation per 10-50 s [52]. For hepatocytes, it was found that 1 oscillation lasted from 20 to 240 s, depending on the IP_3_ concentration [53]. Let us analyze in more detail how cytosolic Ca^2+^ concentration oscillations arise.

### 3.5 Initial step: Ca^2+^ entering into the cytosol through the PM and pumping it out of the cytosol into the SR

As a consequence of the action of the signaling substance, the calcium channels on the PM are opened (the maximum number of all calcium channels located on the PM (0.016 %) opens at approximately 0.00026 s) and Ca^2+^ begins to enter the cytosol. The maximum Ca^2+^ flux into the cytosol through the calcium channels of PM *J*_*p,PM*_ (as well as *J*_*PM*_) is observed when the maximum number of calcium channels of PM are opened (Fig. 4, curve 2).

**Fig. 4.**
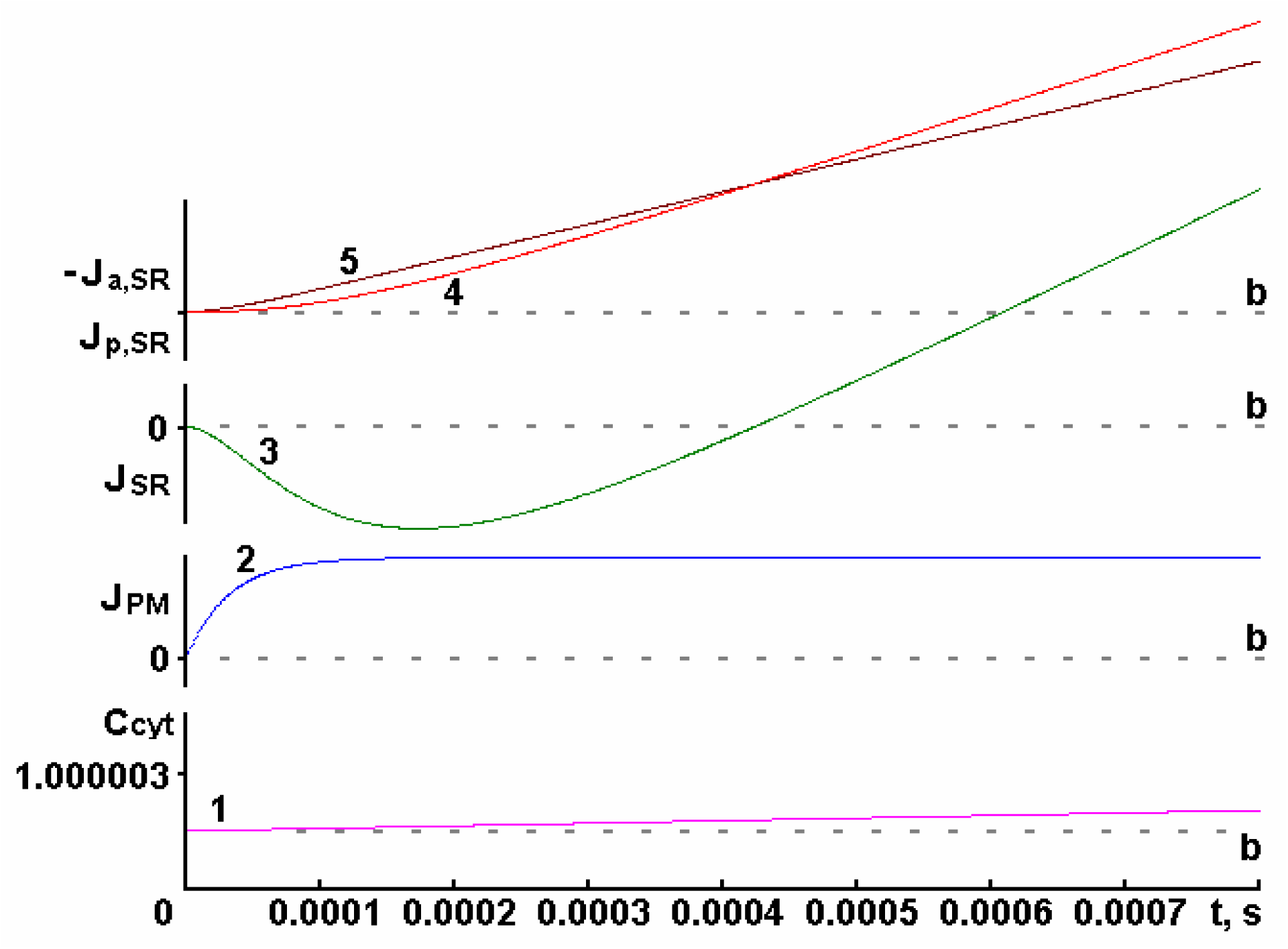
Dependences of Ca^2+^ concentration in cytosol *C*_*cyt*_ = [*Ca*^*2+*^]_*c*_ */* [*Ca*^*2+*^]_*c,b*_ (curve 1) on time *t*, as well as changes of total Ca^2+^ flux through PM *J*_*PM*_ (curve 2), passive Ca^2+^ flux from SR *J*_*p,SR*_ (curve 4), active Ca^2+^ flux into SR *J*_*a,SR*_ (curve 5) and total Ca^2+^ flux through the SR membrane *J*_*SR*_ *= J*_*p,SR*_ + *J*_*a,SR*_ (curve 3) in time (in the initial period of the agonist stimulation process) at *μ*_*SR*_ = 2000, [*A*]_0_ = 10 nM, [*F*]_0_ = 30 nM. The values of other parameters are given in Table 1. The corresponding basal levels are marked with the letter ***b***.

At the same time, additional Ca^2+^ entering the cytosol from the extracellular space begins to be pumped out of the cytosol into the SR (Fig. 4, curve 3), but this lasts only about 0.0004 s. The maximum rate of pumping into the SR (i.e., the maximum *J*_*SR*_) is reached at 0.00017 s.

The phenomenon of Ca^2+^ concentration increase in SR before the calcium-induced calcium release was noted in [54].

### 3.6 Calcium-induced calcium release from SR

An elevated concentration of Ca^2+^ in the cytosol causes an increase in the rate of the calcium pump of SR, which, together with the calcium pump of the PM, begins to pump out of the cytosol the Ca^2+^ entering from the extracellular space. However, as can be seen in Fig. 4, because of the uncompensated passive and active transport from SR, after 0.0004 second there is an increasing flux of Ca^2+^ into the cytosol from the SR, which accelerates the accumulation of Ca^2+^ in the cytosol. This indicates that the calcium pumps of PM and SR do not have enough time to pump the cytosolic Ca^2+^ that enters from the extracellular space and SR. As a result, excess Ca^2+^ opens additional calcium channels of SR (compared to the number of open channels in the steady state), which begin to rapidly pass Ca^2+^ from the SR into the cytosol.

Starting at 0.00043 s (Fig. 4, curve 3), the Ca^2+^ flux from the SR into the cytosol (*J*_*p,SR*_) increases up to 1.389 s (Fig. 5, curve 3), when at its maximum it exceeds the basal level by 2.07 times. The Ca^2+^ concentration in the cytosol increases rapidly by several times (in the given case, by a factor of 6.8 (Fig. 5, curve 1) at the moment of reaching the maximum Ca^2+^ concentration in the cytosol (1.54 s after the beginning of the agonist stimulation process).

**Fig. 5.**
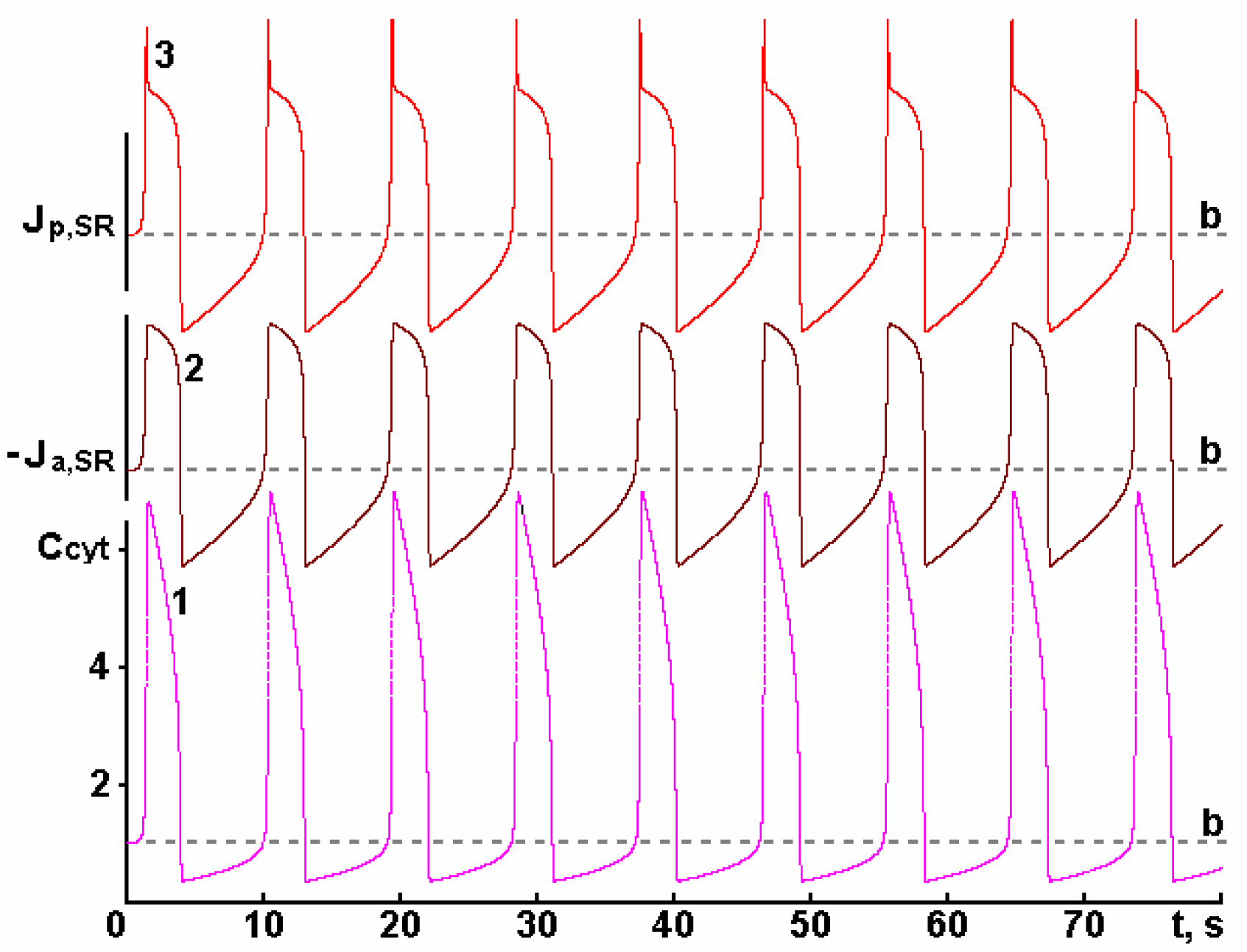
Dependences of Ca^2+^ concentration in cytosol *C*_*cyt*_ = [*Ca*^*2+*^]_*c*_ */* [*Ca*^*2+*^]_*c,b*_ (curve 1) on time *t*, as well as changes of active Ca^2+^ flux in SR *J*_*a,SR*_ (curve 2) and passive Ca^2+^ flux from SR *J*_*p,SR*_ (curve 3) in time at *μ*_*SR*_ = 2000, [*A*]_0_ = 10 nM, [*F*]_0_ = 30 nM. The values of other parameters are given in Table 1. The corresponding basal levels are marked with the letter ***b***.

It was surprising that the maximum Ca^2+^ flux from SR to cytosol (*J*_*p,SR*_) did not coincide in time with the moment of opening of the maximum number of calcium channels on the SR membrane (1.389 and 1.54 s, respectively). This is due to the fact that the magnitude of this flux depends not only on the number of open channels, but also on the difference in Ca^2+^ concentrations between the SR and the cytosol. If the number of open calcium channels increases, the concentration gradient decreases and, as a result, the maximum flux from SR is observed 0.151 s earlier than the moment of opening of the maximum number of calcium channels of SR.

It remains unclear to researchers why the Ca^2+^ entry into the cell from the extracellular space is not in phase with the Ca^2+^ release from SR [55]. Our calculations show that the maximum rate of Ca^2+^ entry into the cytosol from the extracellular space (*J*_*p,PM*_) is reached very quickly (Fig. 4, curve 2) and is observed at the moment of opening of the maximum number of calcium channels of PM at 0.00026 s. At this time, the Ca^2+^ flux from SR (*J*_*p,SR*_, Fig. 4, curve 4) is still not significantly higher than the basal level, as well as the Ca^2+^ flux from the cytosol into SR (*J*_*a,SR*_, Fig. 4, curve 5).

However, *J*_*a,SR*_ exceeds *J*_*p,SR*_ in magnitude, and Ca^2+^ pumping out of the cytosol into the SR occurs at the time point in question (*J*_*SR*_ is negative, Fig. 4, curve 3). *J*_*p,SR*_ begins to exceed *J*_*a,SR*_ only starting from 0.00043 s (although both fluxes continue to increase) and Ca^2+^ starts to be released from SR into cytosol at an increasing rate, opening more and more calcium channels on the SR membrane. The maximum number of open calcium channels of SR (which directly depends on the cytosolic Ca^2+^ concentration) will be at the time when the maximum Ca^2+^ concentration in the cytosol is reached at 1.54 s (Fig. 5, curve 1). However, Ca^2+^ release from the SR lowers Ca^2+^ concentration in the SR, so the maximum Ca^2+^ flux from the SR (*J*_*p,SR*_) will be observed slightly earlier, at 1.389 s (Fig. 5, curve 3). Note that at 1.54 s, when the maximum number of the calcium channels of SR opens (*RR*_*open*_ = 0.00675, i.e., about 230 calcium channels), the Ca^2+^ flux from SR to cytosol (*J*_*p,SR*_) exceeds 1.75 times the basal level, and the Ca^2+^ concentration in SR is several times below the basal level (in the given case 36 % Ca^2+^ remains in SR (Fig. 3, curve 3) relative to basal level). Thus, as a result of the opening of the calcium channels of SR, not all Ca^2+^ contained in the SR is released, but although quite a significant amount of Ca^2+^ remains in SR, process of the calcium-induced calcium release is terminated.

How the process of calcium-induced calcium release from SR into cytosol is terminated is widely discussed in the literature [7, 24, 27, 43, 45, 56]. After all, it seems that the more Ca^2+^ is in the cytosol, the more calcium channels on the SR membrane open, which leads to the release of even more Ca^2+^ from SR into the cytosol, and it is not clear how this avalanche-like process can be terminated [9]. Three hypotheses have been made about the causes of CICR termination [27]: 1) all Ca^2+^ is released from SR; 2) there is Ca^2+^-dependent inactivation of RyR after Ca^2+^ release from SR, which requires a Ca^2+^ concentration in cytosol of 10 mM; 3) all calcium channels (RyR) are closed simultaneously by statistical chance. However, all these reasons are considered by the authors to be highly unlikely. In this regard, let us analyze in detail how CICR termination occurs.

### 3.7 Ca^2+^ fluxes: some trends

In the basal state, because of the presence of 100 nM Ca^2+^ in the cytosol, about 34 channels of all 34050 are open on the SR membrane. The Ca^2+^ flux from the SR through these channels is fully compensated by the activity of the calcium pumps of SR (*J*_*p,SR,b*_ = *J*_*a,SR,b*_ = 7.065.10^-11^ mol.cm^-2.^s^-1^). The opening of calcium channels of PM by signaling substance leads to Ca^2+^ entry into the cytosol, which unbalances the fluxes established in the basal state: the Ca^2+^ flux from the cytosol into the extracellular space (*J*_*a,PM*_) and into SR (*J*_*a,SR*_) increases due to the increase of cytosolic Ca^2+^ concentration (Fig. 4, curve 5), the Ca^2+^ flux from SR to cytosol (*J*_*p,SR*_) increases (although to a lesser extent than *J*_*a,SR*_) because of the increased Ca^2+^ concentration in SR and the opening of additional calcium channels on SR membrane (Fig. 4, curve 4). However, all these fluxes are changed differently over time, resulting in the complex behavior of the whole system.

However, we can observe that (within our model) the number of open calcium channels on SR membrane (*RR*_*open*_), the number of calcium channels closed by Ca^2+^ on the PM (*R*_*close*_), Ca^2+^ fluxes from the cytosol into the extracellular space (*J*_*a,PM*_ and *J*_*a,PM,t*_, Fig. 6, curve 2) and into the SR (*J*_*a,SR*_ and *J*_*a,SR,t*_, Fig. 5, curve 2), and hence the active Ca^2+^ flux from the cytosol (*J*_*a,Cyt*_, Fig. 7, curve 4) are completely determined by the cytosolic Ca^2+^ concentration (see Table 2).

**Table 2.** Passive and active Ca^2+^ fluxes across the entire plasma membrane area (*J*_*p,PM,t*_ and *J*_*a,PM,t*_, respectively) and sarcoplasmic reticulum membrane area (*J*_*p,SR,t*_ and *J*_*a,SR,t*_, respectively), total Ca^2+^ flux across these membranes (*J*_*PM*_.*S*_*PM*_ and *J*_*SR*_.*S*_*SR*_, respectively), passive, active, and total Ca^2+^ fluxes into the cytosol (*J*_*p,cyt*_, *J*_*a,cyt*_ and *J*_*cyt*_, respectively), and Ca^2+^ concentration changes in the extracellular space, cytosol and in the SR (*D*_*out*_ = [*Ca*^*2+*^]_*o*_ *–* [*Ca*^*2+*^]_*o,b*_, *C*_*cyt*_ = [*Ca*^*2+*^]_*c*_ */* [*Ca*^*2+*^]_*c,b*_, *C*_*ret*_ = [*Ca*^*2+*^]_*r*_ */* [*Ca*^*2+*^]_*r,b*_, respectively) at different time points *t* (s). All fluxes are mol.s^-1^ in dimension, and *D*_*out*_, *C*_*cyt*_, *C*_*ret*_ indicate changes in Ca^2+^ concentration relative to the basal level.

| $t, s$ | $D_{out}$ | $C_{cvt}$ | $C_{ret}$ | $J_{p,PM,t}$ | $-J_{a,PM,t}$ | $-J_{a,SR,t}$ | $J_{p,SR,t}$ | $J_{PM}S_{PM}$ | $J_{SR}S_{SR}$ | $-J_{a,cvt}$ | $J_{p,cvt}$ | $J_{cvt}$ |
| --- | --- | --- | --- | --- | --- | --- | --- | --- | --- | --- | --- | --- |
| 0 | 1.0000 | 1.0000 | 1.0000 | 3.996E-19 | 3.996E-19 | 3.996E-17 | 3.996E-17 | 0.000E+00 | 0.000E+00 | 4.036E-17 | 4.036E-17 | 0.00E+00 |
| 0.00017 | 1.0000 | 1.0000 | 1.0000 | 4.002E-19 | 3.996E-19 | 3.996E-17 | 3.996E-17 | 5.770E-22 | -1.420E-24 | 4.036E-17 | 4.036E-17 | 5.75E-22 |
| 0.00026 | 1.0000 | 1.0000 | 1.0000 | 4.002E-19 | 3.996E-19 | 3.996E-17 | 3.996E-17 | 5.774E-22 | -1.178E-24 | 4.036E-17 | 4.036E-17 | 5.76E-22 |
| 0.00043 | 1.0000 | 1.0000 | 1.0000 | 4.002E-19 | 3.996E-19 | 3.996E-17 | 3.996E-17 | 5.770E-22 | 4.120E-26 | 4.036E-17 | 4.036E-17 | 5.77E-22 |
| 0.00046 | 1.0000 | 1.0000 | 1.0000 | 4.002E-19 | 3.996E-19 | 3.996E-17 | 3.996E-17 | 5.770E-22 | 2.793E-25 | 4.036E-17 | 4.036E-17 | 5.77E-22 |
| 0.00077 | 1.0000 | 1.0000 | 1.0000 | 4.002E-19 | 3.996E-19 | 3.996E-17 | 3.996E-17 | 5.770E-22 | 3.035E-24 | 4.036E-17 | 4.036E-17 | 5.80E-22 |
| 0.352 | 1.0000 | 1.00271 | 0.99975 | 4.002E-19 | 4.002E-19 | 4.001E-17 | 4.002E-17 | -1.928E-25 | 1.131E-20 | 4.041E-17 | 4.042E-17 | 1.13E-20 |
| 0.574 | 1.0000 | 1.00999 | 0.99894 | 4.001E-19 | 4.016E-19 | 4.016E-17 | 4.018E-17 | -1.520E-21 | 2.262E-20 | 4.056E-17 | 4.058E-17 | 2.11E-20 |
| 1.376 | 1.00001 | 4.35835 | 0.63385 | 3.999E-19 | 6.501E-19 | 6.498E-17 | 8.167E-17 | -2.502E-19 | 1.669E-17 | 6.564E-17 | 8.207E-17 | 1.64E-17 |
| 1.389 | 1.00001 | 4.85316 | 0.58038 | 3.999E-19 | 6.627E-19 | 6.629E-17 | 8.235E-17 | -2.628E-19 | 1.606E-17 | 6.695E-17 | 8.275E-17 | 1.58E-17 |
| 1.538 | 1.00002 | 6.79810 | 0.36279 | 3.998E-19 | 6.967E-19 | 6.968E-17 | 7.002E-17 | -2.969E-19 | 3.394E-19 | 7.038E-17 | 7.042E-17 | 4.25E-20 |
| 1.540 | 1.00002 | 6.79819 | 0.36263 | 3.998E-19 | 6.967E-19 | 6.968E-17 | 6.996E-17 | -2.969E-19 | 2.828E-19 | 7.038E-17 | 7.036E-17 | -1.41E-20 |
| 1.572 | 1.00002 | 6.78548 | 0.36160 | 3.998E-19 | 6.966E-19 | 6.963E-17 | 6.963E-17 | -2.968E-19 | 3.494E-22 | 7.032E-17 | 7.003E-17 | -2.97E-19 |
| 1.692 | 1.00003 | 6.66170 | 0.36588 | 3.998E-19 | 6.949E-19 | 6.951E-17 | 6.929E-17 | -2.951E-19 | -2.262E-19 | 7.021E-17 | 6.969E-17 | -5.21E-19 |
| 3.878 | 1.00017 | 1.40499 | 0.78323 | 3.999E-19 | 4.668E-19 | 4.668E-17 | 3.976E-17 | -6.688E-20 | -6.923E-18 | 4.715E-17 | 4.016E-17 | -6.99E-18 |
| 3.890 | 1.00017 | 1.20513 | 0.80436 | 3.999E-19 | 4.367E-19 | 4.367E-17 | 3.655E-17 | -3.680E-20 | -7.115E-18 | 4.411E-17 | 3.695E-17 | -7.15E-18 |
| 3.903 | 1.00017 | 0.98967 | 0.82726 | 3.999E-19 | 3.974E-19 | 3.974E-17 | 3.286E-17 | 2.480E-21 | -6.883E-18 | 4.014E-17 | 3.326E-17 | -6.88E-18 |
| 3.975 | 1.00017 | 0.37439 | 0.89482 | 4.000E-19 | 2.177E-19 | 2.177E-17 | 2.086E-17 | 1.823E-19 | -9.106E-19 | 2.199E-17 | 2.126E-17 | -7.28E-19 |
| 4.030 | 1.00017 | 0.34515 | 0.90057 | 4.000E-19 | 2.051E-19 | 2.051E-17 | 2.029E-17 | 1.949E-19 | -2.149E-19 | 2.071E-17 | 2.069E-17 | -2.00E-20 |
| 4.043 | 1.00017 | 0.34486 | 0.90124 | 4.000E-19 | 2.050E-19 | 2.049E-17 | 2.030E-17 | 1.950E-19 | -1.923E-19 | 2.070E-17 | 2.070E-17 | 2.74E-21 |
| 6.901 | 1.00005 | 0.51298 | 1.00001 | 3.998E-19 | 2.710E-19 | 2.710E-17 | 2.700E-17 | 1.288E-19 | -9.615E-20 | 2.737E-17 | 2.740E-17 | 3.26E-20 |
| 8.997 | 1.00000 | 0.72539 | 1.02897 | 3.997E-19 | 3.360E-19 | 3.360E-17 | 3.360E-17 | 6.368E-20 | -9.095E-24 | 3.393E-17 | 3.400E-17 | 6.37E-20 |
| 9.015 | 1.00000 | 0.72810 | 1.02897 | 3.997E-19 | 3.367E-19 | 3.367E-17 | 3.367E-17 | 6.296E-20 | 1.448E-21 | 3.401E-17 | 3.407E-17 | 6.30E-20 |
| 9.922 | 0.99999 | 0.98479 | 1.01054 | 3.997E-19 | 3.966E-19 | 3.965E-17 | 3.997E-17 | 3.120E-21 | 3.167E-19 | 4.005E-17 | 4.037E-17 | 3.20E-19 |
| 9.942 | 0.99999 | 1.00063 | 1.00886 | 3.997E-19 | 3.997E-19 | 3.997E-17 | 4.033E-17 | -8.000E-23 | 3.563E-19 | 4.037E-17 | 4.073E-17 | 3.56E-19 |
| 10.019 | 0.99999 | 1.08350 | 0.99989 | 3.997E-19 | 4.156E-19 | 4.156E-17 | 4.217E-17 | -1.592E-20 | 6.108E-19 | 4.198E-17 | 4.257E-17 | 5.95E-19 |
| 10.252 | 1.00000 | 3.06792 | 0.78391 | 3.996E-19 | 6.028E-19 | 6.029E-17 | 7.426E-17 | -2.032E-19 | 1.397E-17 | 6.090E-17 | 7.466E-17 | 1.38E-17 |
| 10.287 | 1.00000 | 4.41026 | 0.63906 | 3.995E-19 | 6.515E-19 | 6.516E-17 | 8.320E-17 | -2.520E-19 | 1.804E-17 | 6.580E-17 | 8.360E-17 | 1.78E-17 |
| 10.299 | 1.00000 | 4.90676 | 0.58547 | 3.995E-19 | 6.639E-19 | 6.640E-17 | 8.388E-17 | -2.644E-19 | 1.748E-17 | 6.707E-17 | 8.428E-17 | 1.72E-17 |
| 10.319 | 1.00000 | 5.65056 | 0.50500 | 3.995E-19 | 6.790E-19 | 6.793E-17 | 8.224E-17 | -2.795E-19 | 1.431E-17 | 6.861E-17 | 8.264E-17 | 1.40E-17 |
| 10.438 | 1.00001 | 6.96464 | 0.35653 | 3.994E-19 | 6.989E-19 | 6.991E-17 | 7.036E-17 | -2.994E-19 | 4.525E-19 | 7.061E-17 | 7.076E-17 | 1.53E-19 |
| 10.447 | 1.00001 | 6.96637 | 0.35567 | 3.994E-19 | 6.989E-19 | 6.991E-17 | 7.019E-17 | -2.994E-19 | 2.828E-19 | 7.061E-17 | 7.059E-17 | -1.66E-20 |
| 10.479 | 1.00001 | 6.95356 | 0.35464 | 3.994E-19 | 6.987E-19 | 6.985E-17 | 6.985E-17 | -2.993E-19 | 3.388E-22 | 7.055E-17 | 7.025E-17 | -2.99E-19 |
| 12.537 | 1.00016 | 3.91611 | 0.53504 | 3.996E-19 | 6.366E-19 | 6.369E-17 | 6.273E-17 | -2.370E-19 | -9.615E-19 | 6.432E-17 | 6.312E-17 | -1.20E-18 |
| 12.929 | 1.00017 | 1.40477 | 0.78323 | 3.997E-19 | 4.668E-19 | 4.668E-17 | 3.975E-17 | -6.712E-20 | -6.929E-18 | 4.715E-17 | 4.015E-17 | -7.00E-18 |
| 12.941 | 1.00017 | 1.20487 | 0.80437 | 3.997E-19 | 4.366E-19 | 4.367E-17 | 3.655E-17 | -3.696E-20 | -7.115E-18 | 4.410E-17 | 3.695E-17 | -7.15E-18 |
| 12.954 | 1.00017 | 0.98939 | 0.82727 | 3.998E-19 | 3.974E-19 | 3.973E-17 | 3.285E-17 | 2.400E-21 | -6.883E-18 | 4.013E-17 | 3.325E-17 | -6.88E-18 |
| 13.042 | 1.00017 | 0.35593 | 0.89751 | 3.998E-19 | 2.098E-19 | 2.098E-17 | 2.048E-17 | 1.900E-19 | -4.977E-19 | 2.119E-17 | 2.088E-17 | -3.08E-19 |
| 13.081 | 1.00017 | 0.34511 | 0.90055 | 3.998E-19 | 2.050E-19 | 2.050E-17 | 2.029E-17 | 1.947E-19 | -2.149E-19 | 2.071E-17 | 2.069E-17 | -2.02E-20 |
| 13.094 | 1.00017 | 0.34483 | 0.90121 | 3.998E-19 | 2.049E-19 | 2.049E-17 | 2.030E-17 | 1.948E-19 | -1.923E-19 | 2.070E-17 | 2.070E-17 | 2.50E-21 |
| 15.956 | 1.00005 | 0.51301 | 1.00002 | 3.997E-19 | 2.710E-19 | 2.710E-17 | 2.700E-17 | 1.287E-19 | -9.615E-20 | 2.737E-17 | 2.740E-17 | 3.26E-20 |
| 18.054 | 1.00000 | 0.72543 | 1.02897 | 3.996E-19 | 3.360E-19 | 3.360E-17 | 3.360E-17 | 6.360E-20 | 2.072E-23 | 3.394E-17 | 3.400E-17 | 6.36E-20 |
| 18.072 | 1.00000 | 0.72813 | 1.02897 | 3.996E-19 | 3.367E-19 | 3.367E-17 | 3.368E-17 | 6.288E-20 | 5.656E-21 | 3.401E-17 | 3.408E-17 | 6.85E-20 |
| 18.960 | 0.99999 | 0.97056 | 1.01203 | 3.996E-19 | 3.936E-19 | 3.936E-17 | 3.964E-17 | 6.000E-21 | 2.828E-19 | 3.975E-17 | 4.004E-17 | 2.89E-19 |
| 18.980 | 0.99999 | 0.98483 | 1.01053 | 3.996E-19 | 3.966E-19 | 3.965E-17 | 3.997E-17 | 3.040E-21 | 3.167E-19 | 4.005E-17 | 4.037E-17 | 3.20E-19 |
| 19.000 | 0.99999 | 1.00065 | 1.00885 | 3.996E-19 | 3.998E-19 | 3.997E-17 | 4.033E-17 | -1.600E-22 | 3.563E-19 | 4.037E-17 | 4.073E-17 | 3.56E-19 |
| 19.077 | 0.99999 | 1.08345 | 0.99989 | 3.996E-19 | 4.156E-19 | 4.156E-17 | 4.217E-17 | -1.600E-20 | 6.108E-19 | 4.198E-17 | 4.257E-17 | 5.95E-19 |
| 19.179 | 0.99999 | 1.32308 | 0.97352 | 3.996E-19 | 4.552E-19 | 4.552E-17 | 4.726E-17 | -5.560E-20 | 1.736E-18 | 4.597E-17 | 4.766E-17 | 1.68E-18 |
| 19.345 | 1.00000 | 4.40525 | 0.63959 | 3.995E-19 | 6.514E-19 | 6.516E-17 | 8.320E-17 | -2.519E-19 | 1.804E-17 | 6.581E-17 | 8.360E-17 | 1.78E-17 |
| 19.357 | 1.00000 | 4.90185 | 0.58599 | 3.995E-19 | 6.638E-19 | 6.640E-17 | 8.388E-17 | -2.644E-19 | 1.748E-17 | 6.707E-17 | 8.428E-17 | 1.72E-17 |
| 19.377 | 1.00000 | 5.64645 | 0.50543 | 3.994E-19 | 6.790E-19 | 6.787E-17 | 8.224E-17 | -2.795E-19 | 1.437E-17 | 6.855E-17 | 8.264E-17 | 1.41E-17 |
| 19.490 | 1.00001 | 6.96112 | 0.35735 | 3.994E-19 | 6.988E-19 | 6.985E-17 | 7.047E-17 | -2.994E-19 | 6.222E-19 | 7.055E-17 | 7.087E-17 | 3.23E-19 |
| 19.506 | 1.00001 | 6.96612 | 0.35562 | 3.994E-19 | 6.989E-19 | 6.991E-17 | 7.019E-17 | -2.995E-19 | 2.828E-19 | 7.060E-17 | 7.059E-17 | -1.67E-20 |
| 19.537 | 1.00001 | 6.95338 | 0.35465 | 3.994E-19 | 6.987E-19 | 6.985E-17 | 6.985E-17 | -2.994E-19 | 9.990E-22 | 7.055E-17 | 7.025E-17 | -2.99E-19 |
| 21.844 | 1.00017 | 2.91818 | 0.62748 | 3.996E-19 | 5.952E-19 | 5.950E-17 | 5.701E-17 | -1.956E-19 | -2.489E-18 | 6.010E-17 | 5.741E-17 | -2.68E-18 |
| 21.986 | 1.00017 | 1.41241 | 0.78242 | 3.997E-19 | 4.678E-19 | 4.678E-17 | 3.987E-17 | -6.816E-20 | -6.912E-18 | 4.725E-17 | 4.027E-17 | -6.98E-18 |
| 21.998 | 1.00017 | 1.21269 | 0.80354 | 3.997E-19 | 4.379E-19 | 4.379E-17 | 3.668E-17 | -3.824E-20 | -7.115E-18 | 4.423E-17 | 3.708E-17 | -7.15E-18 |
| 22.011 | 1.00017 | 0.99692 | 0.82646 | 3.997E-19 | 3.989E-19 | 3.989E-17 | 3.299E-17 | 8.000E-22 | -6.900E-18 | 4.029E-17 | 3.339E-17 | -6.90E-18 |
| 22.105 | 1.00017 | 0.35249 | 0.89814 | 3.997E-19 | 2.083E-19 | 2.083E-17 | 2.041E-17 | 1.914E-19 | -4.129E-19 | 2.103E-17 | 2.081E-17 | -2.22E-19 |
| 22.138 | 1.00017 | 0.34513 | 0.90052 | 3.997E-19 | 2.050E-19 | 2.050E-17 | 2.029E-17 | 1.946E-19 | -2.149E-19 | 2.071E-17 | 2.069E-17 | -2.03E-20 |
| 22.152 | 1.00017 | 0.34482 | 0.90119 | 3.997E-19 | 2.049E-19 | 2.049E-17 | 2.030E-17 | 1.947E-19 | -1.980E-19 | 2.070E-17 | 2.070E-17 | 2.42E-21 |
| 25.014 | 1.00005 | 0.51300 | 1.00002 | 3.997E-19 | 2.709E-19 | 2.709E-17 | 2.699E-17 | 1.288E-19 | -9.615E-20 | 2.736E-17 | 2.739E-17 | 3.26E-20 |
| 27.113 | 1.00000 | 0.72550 | 1.02897 | 3.996E-19 | 3.360E-19 | 3.360E-17 | 3.360E-17 | 6.360E-20 | 6.026E-23 | 3.394E-17 | 3.400E-17 | 6.36E-20 |
| 27.130 | 1.00000 | 0.72805 | 1.02897 | 3.996E-19 | 3.367E-19 | 3.367E-17 | 3.367E-17 | 6.288E-20 | 1.433E-21 | 3.401E-17 | 3.407E-17 | 6.29E-20 |

**Fig. 6.**
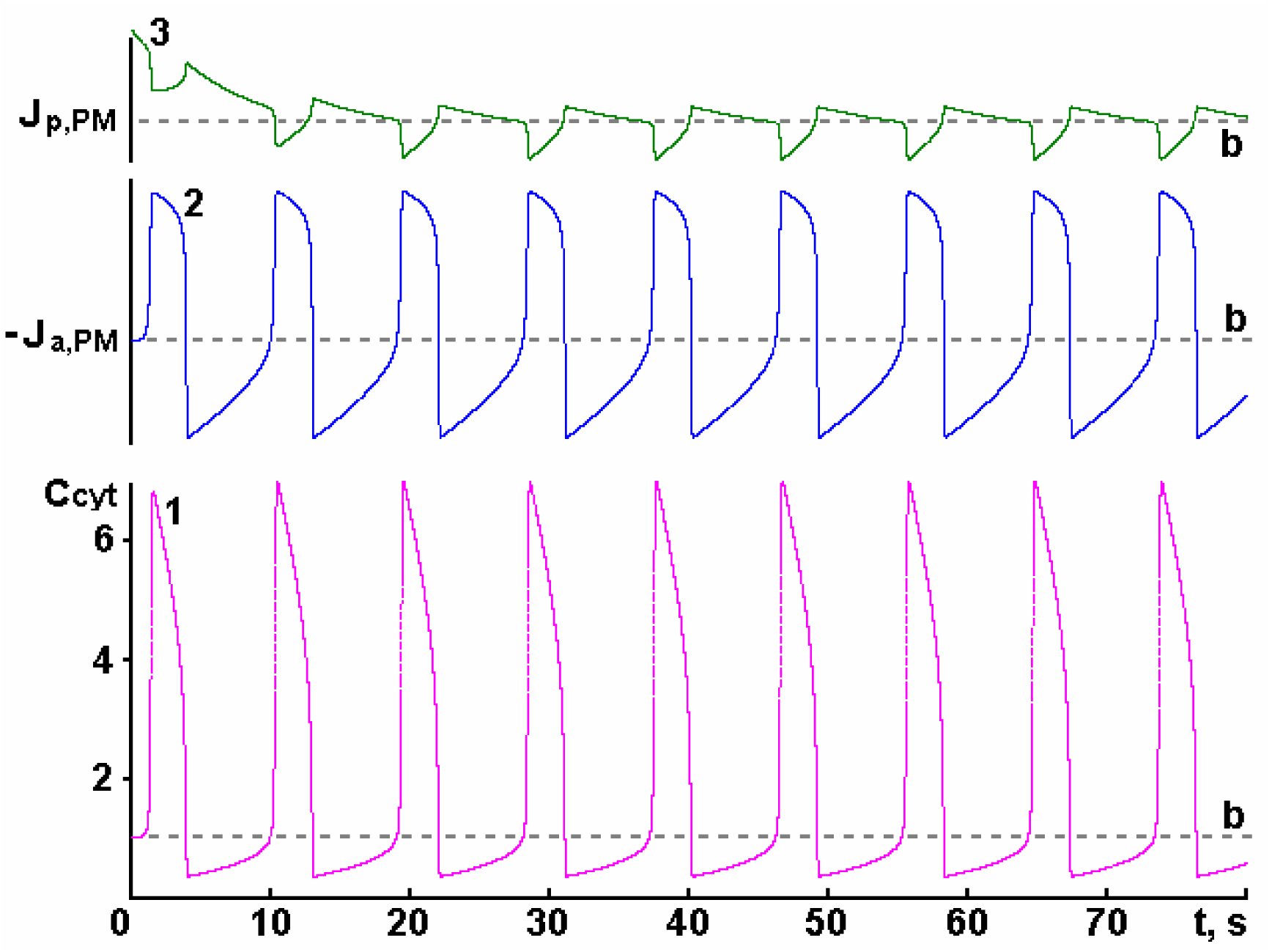
Dependences of Ca^2+^ concentration in cytosol *C*_*cyt*_ = [*Ca*^*2+*^]_*c*_ */* [*Ca*^*2+*^]_*c,b*_ (curve 1) on time *t* as well as changes of active Ca^2+^ flux from cytosol to extracellular space *J*_*a,PM*_ (curve 2) and passive Ca^2+^ flux from extracellular space to cytosol *J*_*p,PM*_ (curve 3) in time at *μ*_*SR*_ = 2000, [*A*]_0_ = 10 nM, [*F*]_0_ = 30 nM. The values of other parameters are given in Table 1. The corresponding basal levels are marked with the letter ***b***.

**Fig. 7.**
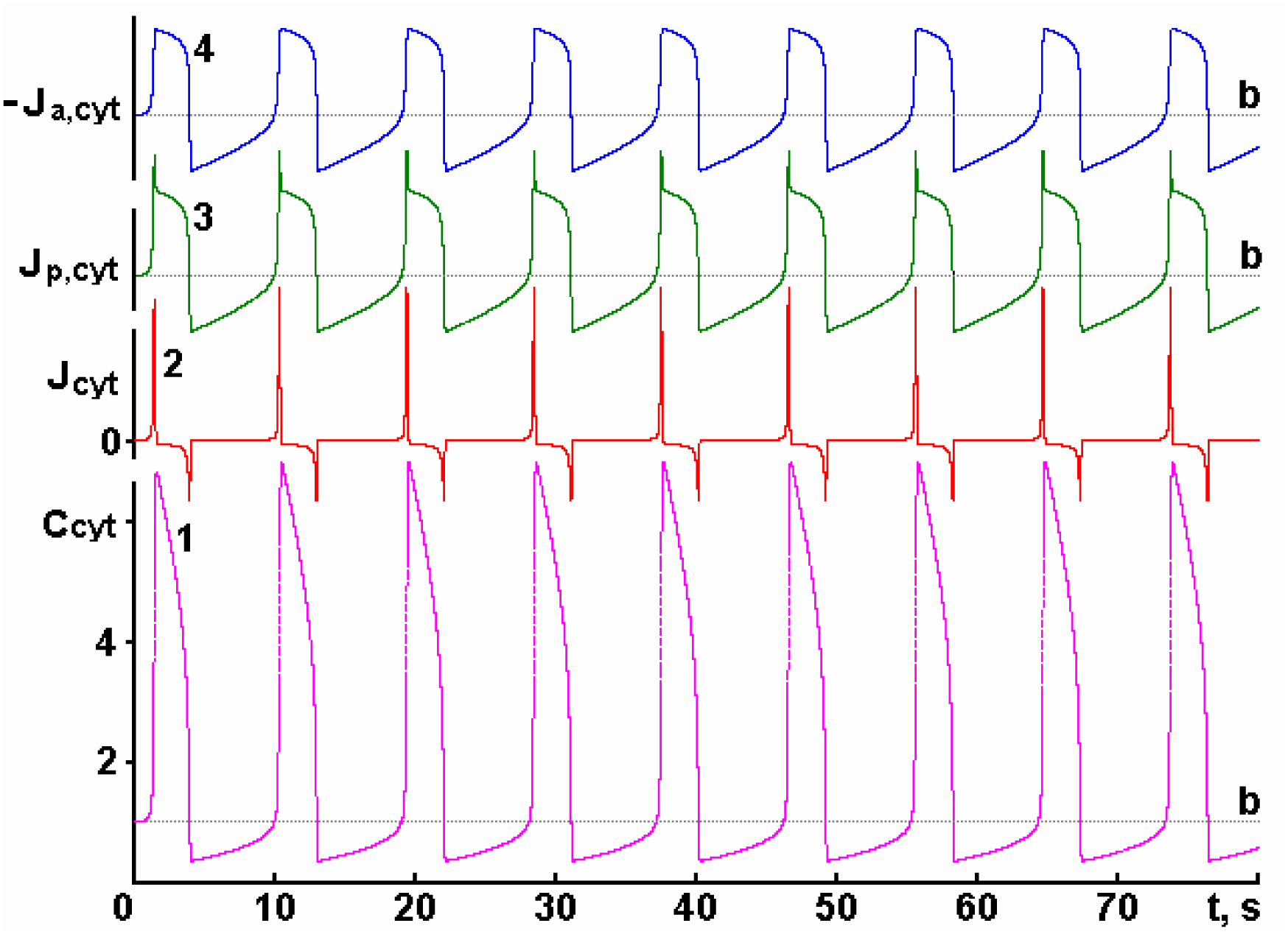
Dependences of Ca^2+^ concentration in cytosol *C*_*cyt*_ = [*Ca*^*2+*^]_*c*_ */* [*Ca*^*2+*^]_*c,b*_ (curve 1) on time *t* as well as changes of total active Ca^2+^ flux from cytosol *J*_*a,Cyt*_ (curve 4), total passive Ca^2+^ flux into cytosol *J*_*p,Cyt*_ (curve 3) and total *J*_*Cyt*_ (curve 2) in time at *μ*_*SR*_ = 2000, [*A*]_0_ = 10 nM, [*F*]_0_ = 30 nM. The values of other parameters are given in Table 1. The corresponding basal levels are marked with the letter ***b***.

Maximum and minimum values of cytosolic Ca^2+^ concentrations (as well as other above-mentioned parameters) are observed at the moments when *J*_*Cyt*_ = 0. Ca^2+^ fluxes into the cytosol from the extracellular space (*J*_*p,PM*_ and *J*_*p,PM,t*_) are predominantly determined by the number of open channels on the PM (*R*_*open*_). The total Ca^2+^ flux through the SR membrane *J*_*SR*_ is zero at the time points when the Ca^2+^ concentration in the SR (*C*_*ret*_) reaches the maximum or minimum value.

In oscillatory processes, Ca^2+^ fluxes from SR to cytosol (*J*_*p,SR*_ and *J*_*p,SR,t*_) and passive Ca^2+^ flux to cytosol (*J*_*p,Cyt*_) change synchronously, which is explained by two orders of magnitude higher values of *J*_*p,SR,t*_ compared to *J*_*p,PM,t*_ (see Table 2).

With the aim of gaining a clearer understanding of how the oscillatory process is carried out, Table 2 presents the numerical values of Ca^2+^ concentrations in the extracellular space, cytosol, and SR, as well as different Ca^2+^ fluxes at the time points when some of them reach the maximum (in red), minimum (in blue), or basal (in pink) level. Based on the data in Table 2, the sequence of events occurring in the cellular system during the oscillatory process can be traced.

### 3.8 How calcium-induced calcium release from the SR is terminated

A rapid elevation of cytosolic Ca^2+^ concentration significantly increases the rate of the calcium pumps of PM and SR: the rate of both increases by about 74.4 % at *t*_*max,1*_ = 1.54 s after the start of stimulation. When the maximum Ca^2+^ concentration in the cytosol is reached (*C*_*cyt,max,1*_ = 6.8), the passive Ca^2+^ flux into the cytosol through the calcium channels of PM and SR (*J*_*p,Cyt*_) and the active Ca^2+^ flux from the cytosol driven by the calcium pumps of PM and SR (*J*_*a,Cyt*_) become equal (*J*_*Cyt*_ = 0) (see Fig. 7). At this time SR is emptied by almost a third (*C*_*ret*_ = 0.36), therefore, although the number of calcium channels open on the SR membrane at this time is maximal, the Ca^2+^ flux from SR to cytosol (*J*_*p,SR*_) already decreases for 0.15 s (and during this time it decreases by 15 % of the maximal level).

The active flux from the cytosol (*J*_*a,Cyt*_) at this time reaches the maximum level and becomes equal in magnitude to the passive flux into the cytosol (*J*_*p,Cyt*_), which, like the *J*_*p,SR*_ flux, decreases as early as 0.15 s (see Fig. 5 and Table 2). As a result *J*_*Cyt*_ = 0 and Ca^2+^ accumulation in cytosol is terminated. Thus, although it appears that Ca^2+^ should be released into the cytosol from the SR at an accelerating rate, in fact, at a certain point in time the contribution of the number of open calcium channels on the SR membrane that can be opened at a given concentration of Ca^2+^ in the cytosol can no longer exceed the effect of the decreasing Ca^2+^ concentration in SR (see equation (29)). This leads to the fact that the passive Ca^2+^ flux into the cytosol (*J*_*p,Cyt*_), after passing through the maximum, begins to decrease (Fig. 7, curve 3).

### 3.9 Ca^2+^ pumping out of cytosol below basal level

Since 1.54 s, when *J*_*Cyt*_ = 0, the process of Ca^2+^ pumping out of the cytosol is started. Note that the Ca^2+^ pumping out of the cytosol by the calcium pumps of the PM and SR is started immediately after begins to enter the cytosol, but, nevertheless, up to 1.54 s the passive Ca^2+^ flux into the cytosol *J*_*p,Cyt*_ exceeded the active Ca^2+^ flux from the cytosol *J*_*a,Cyt*_ (see Table 2). After 1.54 s, both calcium fluxes decrease (*J*_*p,Cyt*_ up to 4.030 s and *J*_*a,Cyt*_ up to 4.043 s), but the passive Ca^2+^ flux into the cytosol decreases more intensively than the active Ca^2+^ flux (Fig. 7, curves 3 and 4, respectively), so that *J*_*Cyt*_ becomes less than zero (Fig. 7, curve 2), i.e., Ca^2+^ is pumped out of the cytosol (the maximum pumping rate is reached at 3.89 s and continues up to 4.043 s, see Table 2). As a result, the cytosolic Ca^2+^ concentration sharply decreases (Fig. 7, curve 1), and this occurs so quickly that at *t*_*min,1*_ = 4.043 s it drops several times below the basal level (approximately 3 times in the analyzed case, *C*_*cyt,min,1*_ = 0.34).

Why does this happen? At the time when the Ca^2+^ concentration in the cytosol reaches the basal level (at approximately 3.903 s), the Ca^2+^ concentration in the extracellular space *C*_*out*_ maximally exceeds the basal level (Fig. 3, curve 2), while in SR the Ca^2+^ concentration (Fig. 3, curve 3) has not yet reached the basal level (it will be reached only at 6.901 s). At this time, there are driving forces (due to the concentration gradients) for Ca^2+^ transport from the extracellular space to the cytosol (Fig. 8, curve 2, *J*_*PM*_ is positive, even if there are no calcium channels open with the signal substance on the PM; see equation (27)) and from the cytosol to SR (*J*_*SR*_ is negative, Fig. 8, curve 3), because the Ca^2+^ flux from the cytosol to SR (*J*_*a,SR*_) is equal to the basal level, according to equation (30), and the Ca^2+^ flux from SR to cytosol (*J*_*p,SR*_) is less than the basal level, according to equation (29) (see Table 2). However, because the Ca^2+^ flux from the cytosol to the SR (*J*_*SR*_) is greater than the Ca^2+^ flux from the extracellular space to the cytosol (*J*_*PM*_) (at 3.903 seconds, see Table 2), the Ca^2+^ concentration in the cytosol decreases below the basal level. Only at 4.043 s *J*_*SR*_ = *J*_*PM*_ and *J*_*Cyt*_ = 0, resulting in the termination of further decrease of cytosolic Ca^2+^ concentration. However, the above-mentioned driving forces did not disappear, but only changed their intensity.

**Fig. 8.**
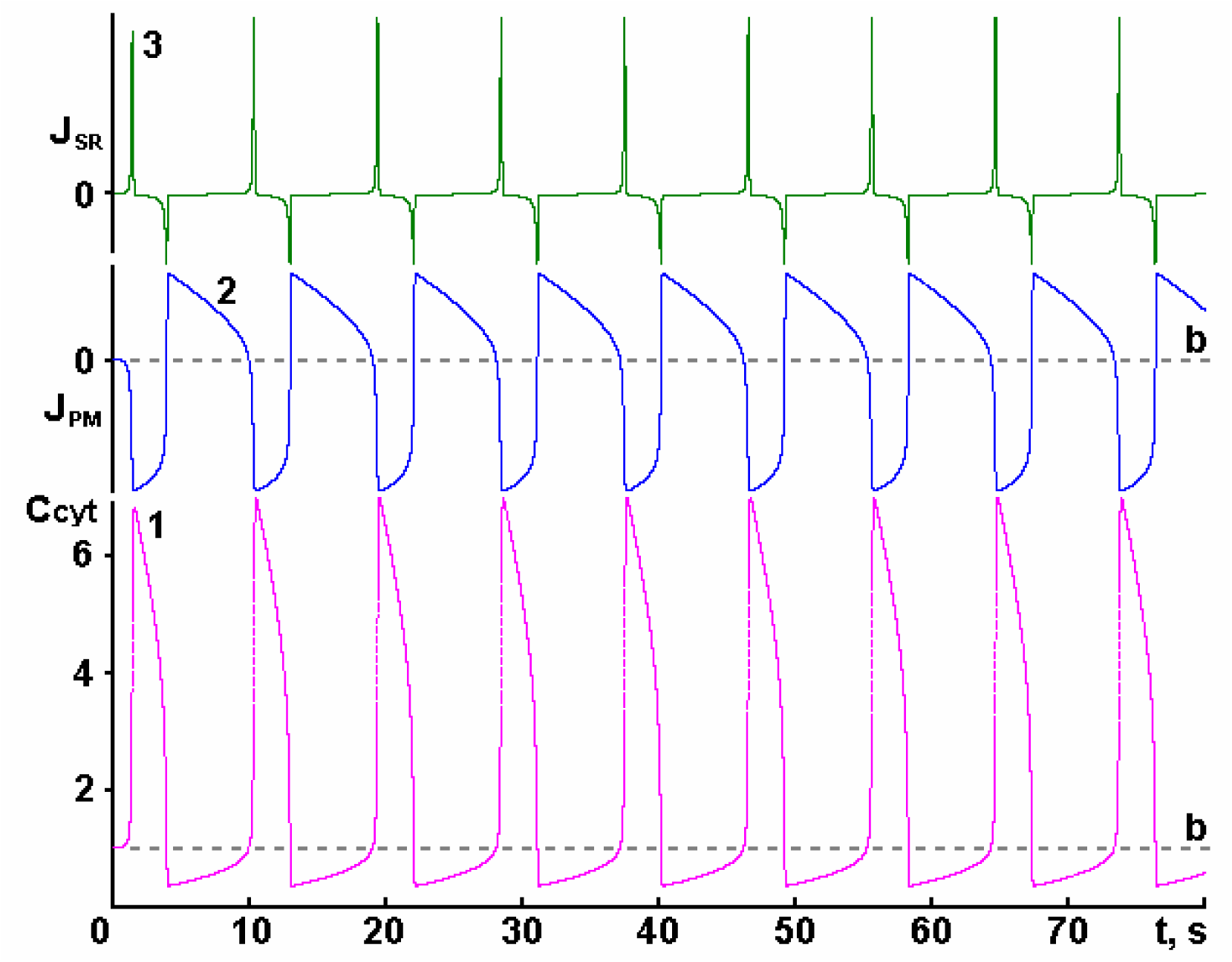
Dependences of Ca^2+^ concentration in cytosol *C*_*cyt*_ = [*Ca*^*2+*^]_*c*_ */* [*Ca*^*2+*^]_*c,b*_ (curve 1) on time *t* as well as the change in total Ca^2+^ flux through PM *J*_*PM*_ (curve 2) and total Ca^2+^ flux through SR membrane *J*_*SR*_ (curve 3) in time at *μ*_*SR*_ = 2000, [*A*]_0_ = 10 nM, [*F*]_0_ = 30 nM. The values of other parameters are given in Table 1. The corresponding basal levels are marked with the letter ***b***.

As a result of the increase of Ca^2+^ concentration in the SR (approaching but not exceeding the basal level), the Ca^2+^ flux from the cytosol to the SR (*J*_*SR*_) becomes less than the Ca^2+^ flux from the extracellular space to the cytosol (*J*_*PM*_) (although the Ca^2+^ concentration in the extracellular space decreased, also approaching the basal level). Consequently, slow increase of Ca^2+^ concentration in the cytosol is started coupled with an increase of Ca^2+^ concentration in the SR (Fig. 3, curves 1 and 3, respectively).

Thus, due to the action of the above-mentioned driving forces, the SR refilling at the expense of extracellular Ca^2+^ is started in the cell system. Note that the low intensity of these forces (caused by relatively small deviations of Ca^2+^ concentrations in the cytosol, extracellular space, and SR from the basal values) leads to a very slow increase of the cytosolic Ca^2+^ concentration.

### 3.10 SR refilling: analogy with the SOCE process

Since the time of 4.043 s, when the *J*_*Cyt*_ flux becomes positive and Ca^2+^ predominantly enters the cytosol (see Table 2), the next step of the process begins: the Ca^2+^ concentration in the cytosol, SR, and extracellular space slowly approaches the basal level (see Fig. 3). At this step, the Ca^2+^ concentration in the cytosol slowly increases, remaining below the basal level (the basal level is reached only at 9.942 s), as well as the Ca^2+^ concentration in the SR (reaching the basal level already at 6.901 s). Afterwards it begins to exceed the basal level and the maximum excess is observed at 8.997 s (*C*_*ret*_ = 1.029), i.e., before the cytosolic Ca^2+^ concentration reaches the basal level. At the same time, the Ca^2+^ concentration in the extracellular space is constantly decreasing (the basal level is reached at 9.015 s, and the minimum level at 9.942 s) and, in general, the situation at this step can be interpreted as functioning of SOCs – at very low cytosolic Ca^2+^ concentration, not exceeding basal level, SR is filled by Ca^2+^ entering from extracellular space [2, 7, 8, 12, 22, 55, 56].

As can be seen, the effect of the SR refilling by extracellular Ca^2+^ can be explained without the use of additional intracellular components. There is no need to consider the existence of a direct connection of the SR to the extracellular space [8, 12, 55] or a special intracellular messenger [8, 12, 55] or G-protein [12, 55], or protein interactions between receptors of calcium channel of SR and calcium channels of PM [12, 55], or vesicles with proteins of SOCs [12], or differences in cytosolic Ca^2+^ concentrations in the volume and near the PM [12]. All the components listed above are absent from our model. Observed effect is purely kinetic, arising as a result of an excess of Ca^2+^ in the extracellular space compared to the basal state, while there is a Ca^2+^ deficiency in the SR at the same time (pumped out of the cytosol into the extracellular space during CICR) – thus generating driving forces to refill the SR at the expense of extracellular Ca^2+^.

### 3.11 The next massive Ca^2+^ release from the SR into the cytosol

At 9.942 s, when Ca^2+^ concentration in the cytosol reaches its basal level (*C*_*cyt*_ = 1), the extracellular Ca^2+^ concentration is at its minimum, i.e., below the basal level, while the Ca^2+^ concentration in the SR still exceeds the basal level (*C*_*ret*_ = 1.0089) (see Fig. 3 and Table 2). At this time, the situation is the exact opposite of the one observed at 3.903 s: there are driving forces for Ca^2+^ transport from the cytosol to the extracellular space (*J*_*PM*_ becomes negative at 9.942 s and reaches its maximum value (in absolute value) at 10.447 s) as well as from SR to cytosol (*J*_*SR*_ is positive, because the Ca^2+^ flux from cytosol to SR *J*_*a,SR*_ is equal to basal, while the Ca^2+^ flux from SR to cytosol *J*_*p,SR*_ is greater than basal level). Ca^2+^ flux from the SR to the cytosol (*J*_*SR*_) increases approximately 50-fold from 9.942 to 10.287 s (when the *J*_*SR*_ reaches its maximum value) (Fig. 8, curve 3). In this way, the next massive Ca^2+^ release from the SR into the cytosol occurs. Ca^2+^ release continues until 10.447 s (see Table 2), when the Ca^2+^ concentration in SR becomes practically minimal (*C*_*ret*_ = 0.355). As a result of this Ca^2+^ release, the Ca^2+^ concentration in the extracellular space increases, overcoming the basal level at 10.319 s, and the Ca^2+^ concentration in the cytosol reaches a maximum at *t*_*max,2*_ = 10.447 s (*C*_*cyt,max,2*_ = 6.966) (see Fig. 3). At this time *J*_*a,Cyt*_ reaches its maximum value, but *J*_*p,Cyt*_ passed its maximum almost 0.15 s earlier, and these fluxes become equal: as a result, *J*_*Cyt*_ = 0 and Ca^2+^ accumulation in the cytosol is terminated (see Fig. 7). This is the end of the cycle of Ca^2+^ release from SR into the cytosol and a new cycle of Ca^2+^ pumping out of the cytosol is started.

The periodic oscillation mode is possible only when there are three compartments. When two compartments exist, this mode is not observed [33]. Periodic oscillations arise because in the cellular system Ca^2+^ concentration cannot reach the basal level in the three compartments simultaneously. Instead, Ca^2+^ constantly “flows” from the extracellular space into the sarcoplasmic reticulum and back through the cytosol, causing oscillations not only of cytosolic Ca^2+^, but also of extracellular Ca^2+^ and Ca^2+^concentration in the SR. The oscillatory mode involves all compartments.

### 3.12 Maintaining calcium balance during cell stimulation

It can be seen that tuning through the basal state allows the cellular system to return exactly to the initial state after finishing the agonist stimulation. This permits us to answer the question raised in [8]: how is it possible that the amount of Ca^2+^ passing through the SOCs is exactly equal to the amount of Ca^2+^ received by the SR, so that from extracellular space enters the cell exactly the same amount of Ca^2+^ that then leaves it during agonist stimulation? This is now obvious: due to the return of the cellular system strictly to its initial state the above-mentioned Ca^2+^ balance is maintained. If we assume that in the SOCE process the Ca^2+^ concentration in the cytosol does not change and is equal to the basal one, then the amount of Ca^2+^ entering the cell from the extracellular space (lowering the Ca^2+^ concentration in the extracellular space from the excess level to the basal one) will be exactly equal to the amount of Ca^2+^ that will be received by the SR (due to increase of Ca^2+^ concentration in the SR from the reduced level to the basal one), because Ca^2+^ does not disappear anywhere, but is only redistributed between the extracellular space, cytosol and SR. Actually, as it was mentioned above, the possibility of Ca^2+^ redistribution between these three compartments creates the possibility of cytosolic Ca^2+^ oscillations: after removing any of these three compartments (i.e., removing one of the membranes) the cell system cannot switch into oscillatory mode [33]. Therefore, we can agree with [53] that the oscillatory mode must be accompanied by oscillations of the extracellular Ca^2+^ concentration and with [2] that Ca^2+^ entry through CRAC channels is necessary to maintain Ca^2+^ oscillations in the cytosol.

### 3.13 A small number of calcium channels on the plasma membrane opened by the agonist are closed by cytosolic Ca^2+^

Analysis of the model shows that there is no special need to involve the idea that cytosolic Ca^2+^ must close the calcium channels of the PM to terminate Ca^2+^ flow from the extracellular space. Indeed, the results show that despite rather high values of the rate constant of Ca^2+^ binding to the PM, the percentage of closed channels of the PM by cytosolic Ca^2+^ is negligible (in the basal state *R*_*close*_ *=* 0.003322, i.e. only 0.33 % of the calcium channels are closed, and the maximum number of closed channels is observed at *t*_*max,2*_ = 10.447 s – 2.27 % of all those opened with the signal substance).

### 3.14 The agonist binds with a small number of receptors

It is also important to emphasize that in spite of high values of the rate constants of signal substance binding with the receptors of calcium channel on PM and cytosolic Ca^2+^ with the receptors of calcium channel on SR membrane, the number of open channels on the PM and SR membrane remains rather insignificant. As shown earlier, in the basal state (i.e., at a cytosolic Ca^2+^ concentration of 100 nM), 34 or 0.0999 % of 34050 calcium channels of SR are open. The maximum number of SR channels open at *t*_*max,2*_ = 10.447 s – *RR*_*open*_ = 0.0069, i.e. 236 or 0.69 %. This value is in good agreement with the literature, which suggests that the number of ryanodine receptors that bind with Ca^2+^ (when Ca^2+^ flux from the SR is maximal) is 1-2 % [3]. The maximum number of open calcium channels on PM at [*A*]_0_ = 10 nM is 773 out of approximately 4.816.10^6^ located on the PM, which is approximately 0.016 % (*R*_*open*_ = 0.00016). At an agonist concentration of 1 μM, approximately 1.5 % of the Ca^2+^ channels on the PM may be open. In the literature, it is reported that the maximum response is usually observed when the agonist binds with 2-3 % of the PM receptors [35, 40]. Despite such a small number of opening Ca^2+^ channels, the process of Ca^2+^ accumulation in the cytosol proceeds comparable with the experiment.

### 3.15 How many calcium cations are needed to start CICR and can so much Ca^2+^ from a calcium-free solution enter the cytosol?

Let us return to the process of calcium-induced calcium release and analyze the known facts taking into account the results obtained in the study of our model. It is known that when a smooth muscle cell is stimulated by agonists in a calcium-free environment, Ca^2+^ release from SR is observed [8]. Based on this fact, it can be concluded that calcium channels on SR membrane are not opened by Ca^2+^ (which as it is believed is absent in the external solution), but indirectly by IP_3_.

As our model shows, the opening of Ca^2+^ channels on SR membrane can also be performed by Ca^2+^, because it requires only a few, tens or hundreds of free calcium cations (as shown above the whole process of calcium-induced calcium release within our model is performed under conditions when only 202 Ca^2+^ channels open on the SR membrane in addition to the basal level, i.e. only 202 calcium cations bind with the RyR when the maximum Ca^2+^ concentration in cytosol is reached) that always exist in the external solution (due to the existence of thermodynamic equilibrium).

It is difficult to calculate the amount of calcium cations required to initiate the CICR process because Ca^2+^ simultaneously enters the cytosol from the extracellular space and SR and at the same time is pumped out of the cytosol into SR and the extracellular space. However, if we take the Ca^2+^ flux into the cytosol from the extracellular space, which changes relatively little during the process, equal to approximately *J*_*p,PM*_ = 4.10^-19^ mol/s, and calculate how many calcium cations will enter the cytosol during 0.0004 s (approximately until the Ca^2+^ flux from SR to cytosol begins to exceed the Ca^2+^ flux from cytosol to SR (see Fig. 4), that is, when *J*_*SR*_ flux becomes positive), we get 96 calcium cations. Thus, an increase in the Ca^2+^ content of the cytosol with only 96 calcium cations is enough to provoke the process of Ca^2+^ release from the SR (naturally, with the parameters we chose for our calculations).

It is known that Krebs solution without the addition of Ca^2+^or so called “calcium-free” medium actually contains about 1.10^-5^ M Ca^2+^ and addition of 0.1 mM EGTA lowers Ca^2+^ concentration to about 1.10^-8^ M [49]. It can be calculated that at a concentration of 1.10^-5^ M Ca^2+^ there are about 24 million calcium cations in the extracellular space in a free state, and at a concentration of 1.10^-8^ M there are 2.4.10^4^ (if the volume of extracellular space is 4 picoliters, as in the framework of our model). Therefore, one cannot assume that the calcium-free medium is completely free of calcium ions, and our model shows that much fewer calcium cations than are contained in the calcium-free medium may be required to initiate the calcium-induced calcium release from the SR.

### 3.16 What studies can be carried out using the developed calcium-induced calcium release model?

In order to model how the cytosolic Ca^2+^ signal depends on the agonist *in vivo*, it is necessary to consider the presence of an agonist-degrading enzyme in the cell system, which allows it to return to its initial state after termination of stimulation. The above results refer to this type of study. As can be seen, an increase in the conductance gain coefficient of the calcium channel of SR favors the appearance of an oscillatory mode.

At small values of the conductance gain coefficient of calcium channel of SR (*μ*_*SR*_ = 50) in the cell system the oscillatory mode is not realized (see Fig. 1), and the amplitude of the calcium transient in the cytosol increases with increasing agonist concentration. At low agonist concentrations, the change in Ca^2+^ concentration in the cytosol is almost undetectable. For example, at an agonist concentration of 0.01 μM ([*F*]_0_ = 30 HM, *μ*_*SR*_ = 50) the cytosolic Ca^2+^ concentration changes by only 0.1 %. At higher agonist concentrations (up to 1 μM), the amplitude of calcium transient in the cytosol increases markedly, and SR in this case accumulates part of Ca^2+^ entering the cytosol (see Fig. 1, curve 7). At even higher concentrations of the agonist Ca^2+^ begins to leave the SR, initially in an amount less than was previously pumped into it (in the range of agonist concentrations 1-20 μM, see Fig. 1, curve 8), and then the amount of Ca^2+^ leaving the SR begins to exceed the amount of Ca^2+^ previously pumped (in the range of agonist concentrations from 20 μM to 1 mM, see Fig. 1, curve 9). Note that the maximum degree of sarcoplasmic reticulum emptying at an agonist concentration of 1 mM is 83 %.

As the agonist concentration increases further, the behavior of the SR becomes even more complex. During the period of increasing cytosolic Ca^2+^ concentration, the SR first pumps Ca^2+^ from the cytosol, then calcium-induced calcium release occurs, after which the slow pumping of cytosolic Ca^2+^ into the SR begins again. After the Ca^2+^ concentration in the cytosol begins to decrease, its content in the SR also begins to decrease slowly at first, and then a rapid refilling of the SR with Ca^2+^ occurs, so rapidly that the Ca^2+^ content in the SR exceeds the basal level. The Ca^2+^ content in the SR then begins to decrease until it reaches the basal level (at the same time the cytosolic Ca^2+^ concentration also reaches the basal level). On the contrary, at high values of the conductance gain coefficient of calcium channel of SR (*μ*_*SR*_ = 2000), only the oscillatory mode is realized. This was tested in the range of agonist concentrations from 0.01 nM to 100 mM.

To model *in vitro* studies, the effect of the agonist-degrading enzyme can be ignored. Under such conditions it is possible to study not only the effect of the agonist on Ca^2+^ distribution in the cell system but also the effect of inhibition or activation of calcium pumps of the plasma membrane and SR [57]. In the absence of an agonist-degrading enzyme, the cell system will not return to the initial (basal) state but will switch to a new steady state different from the basal one. Note that in many aspects the action of the agonist is very similar to the action of inhibitors of calcium pumps of PM.

As the agonist concentration or the degree of inhibition of the plasma membrane calcium pumps increases, the following shapes of the time dependence of the Ca^2+^ concentration in the cytosol can be observed:

1. the level of Ca^2+^ concentration in the cytosol remains virtually unchanged (at very low agonist concentrations).
2. the Ca^2+^ concentration in the cytosol slowly and monotonically increases until it reaches a new steady-state level (the SR accumulates some Ca^2+^ entering the cytosol, and no calcium-induced calcium release occurs).
3. a maximum of cytosolic Ca^2+^ with small amplitude appears, after which its concentration decreases to a new stationary level. There is no massive Ca^2+^ release from the SR. In some cases (at relatively low agonist concentrations or low *μ*_*SR*_) some Ca^2+^ entering the cytosol may be accumulated by the SR and in other cases (at relatively high agonist concentrations or high *μ*_*SR*_) a weak Ca^2+^ flux into the cytosol from the SR may be observed.
4. there is a massive release of Ca^2+^ from the SR (CICR). The amplitude of the calcium transient increases significantly. The new steady-state level of Ca^2+^ concentration in the cytosol is determined by both the agonist concentration and the degree of inhibition of the calcium pump of PM: the greater these parameters, the higher the new steady state level.
5. damped oscillations of Ca^2+^ concentration in the cytosol appear. The large amplitude of the first maximum is a consequence of calcium-induced calcium release from the SR. The amplitude of the second (and subsequent) maxima is small at first but increases as the concentration of the agonist or the degree of inhibition of the calcium pump of PM increases. Note also that damped oscillations may also appear with relatively small Ca^2+^ influx into the cytosol from the SR, but in this case the oscillations may remain almost undetectable (see point 3).
6. a mode of periodic oscillations of Ca^2+^ concentration in the cytosol is established. With the increase of agonist concentration, the amplitude of the second and subsequent maxima first increases and then begins to decrease.
7. a decrease in the amplitude of the second and subsequent maxima is the reason why periodic oscillations turn into damped ones.
8. the damped oscillations are terminated. The cytosolic calcium signal again takes the form of a transient, and the new stationary level of cytosolic Ca^2+^ concentration appears to be significantly higher than the basal level (by orders of magnitude).
9. the Ca^2+^ concentration in the cytosol after a sharp increase as a result of calcium-induced calcium release from the SR remains almost at the same level, neither significantly decreasing nor increasing. The cytosolic calcium signal loses its transient form. At this time, the step of Ca^2+^ pumping into the SR from the cytosol is absent, although a huge amount of Ca^2+^ enters the cytosol from the extracellular space.
10. the Ca^2+^ concentration in the cytosol after a sharp increase as a result of calcium-induced calcium release from the SR slowly and monotonically increases until it reaches a new stationary level. In this region, the maximum possible rate of the calcium pump of PM is less than the rate of Ca^2+^ entry into the cytosol from the extracellular space, so its concentration in the cytosol increases until the rate of Ca^2+^ entry into the cytosol from the extracellular space becomes equal to the maximum rate of the calcium pump of PM (the decrease in the rate of Ca^2+^ entry into the cytosol is achieved by significantly increasing its concentration in the cytosol, which reduces the Ca^2+^ concentration gradient between the cytosol and the extracellular space).

Note that not all of the above shapes of the cytosolic Ca^2+^ concentration time dependences can be observed in each specific case. For example, as was shown above, damped and periodic oscillations are not observed at small values of *μ*_*SR*_, and only periodic oscillations are observed at its large values in the range of agonist concentrations from 0.01 nM to 100 mM.

A similar situation, when in some types of smooth muscle cells the agonist induces cytosolic Ca^2+^ oscillations, while in other types only transient Ca^2+^ signal is observed, is described, for example, in [58].

### 3.17 More about SR and SOC properties

The results of the study of our model show that the SR is never completely emptied (this is also noted in [2, 3, 8, 24]), although it seems that the massive Ca^2+^ release from the SR should continue until the SR is completely emptied [8, 9, 27]. In the oscillatory mode, the maximum degree of SR emptying is 64.5 % (*C*_*ret*_ = 0.355 at time at *t*_*max,2*_ = 10.447 s). This is quite close to the values reported in the literature, in particular, in the article [2] for pancreatic smooth muscle cells it is indicated that SR is emptied by 40 % during one spike.

Sometimes the Ca^2+^ release from SR is not accompanied by the opening of SOCs [2]. It can be seen in Fig. 1 that in the case when the conductance gain coefficient of calcium channels of SR is relatively low, there is no SOC-like effect. After Ca^2+^ release from the SR has occurred, Ca^2+^ is pumped out of the cytosol into the SR and into the extracellular space, that is, at the time when the SR is filling, the Ca^2+^ flow through the PM *J*_*PM*_ is mainly directed from the cytosol into the extracellular space rather than into the cytosol. Therefore, we cannot say that Ca^2+^ transferring the SR arriving from the extracellular space (via SOCs).

One of the features of the SOCs is that if the stimulating action of a signaling substance is terminated, these channels remain open until the Ca^2+^ stores in SR are completely refilled [2]. A similar effect is indeed observed in the study of our model: as can be seen from Fig. 3, at the time when the seventh maximum of Ca^2+^ concentration in the cytosol is observed, the signal substance concentration (curve 4) is only 0.003 % of the initial one. We can assume that the action of the signaling substance is practically absent, however, even after this moment the processes of SR refilling and simultaneous Ca^2+^ entry into the cytosol from the extracellular space continue.

### 3.18 Whether SOCs are needed for SR refilling: modeling an experimental protocol

Above, we have drawn an analogy between the SR refilling with simultaneous entry of Ca^2+^ into the cytosol from the extracellular space, which we observed within the developed model in the oscillatory mode, with the SOCE process observed in experimental studies. However, these studies are performed under specific conditions.

To identify the calcium channels of PM that open in response to SR emptying and allow SR refilling, an experimental protocol is used where thapsigargin (TG), an inhibitor of calcium pump of SR, is applied in the presence of a calcium-free extracellular solution, which causes SR emptying. Then returns the Ca^2+^ concentration in the extracellular solution to the basal level and observes an increase in Ca^2+^ concentration in the cytosol, which is interpreted as a sign of SOCs opening and Ca^2+^ entry through them into the cytosol from the extracellular solution.

Within the developed model, we carried out studies simulating the protocol described above (Fig. 9). The results obtained mostly coincide with the experimental data. The effect of thapsigargin was modeled by changing the limiting rate of the calcium pump of SR *V*_*mSR*_ to zero, which corresponds to complete inhibition of this pump. Replacement of the extracellular solution, which contained Ca^2+^ at the basal level, with a calcium-free solution was modeled by a change of Ca^2+^ concentration in the extracellular space from 1.10^-3^ M to 1.10^-8^ M, which corresponds to the Ca^2+^ concentration in the presence of EGTA.

**Fig. 9.**
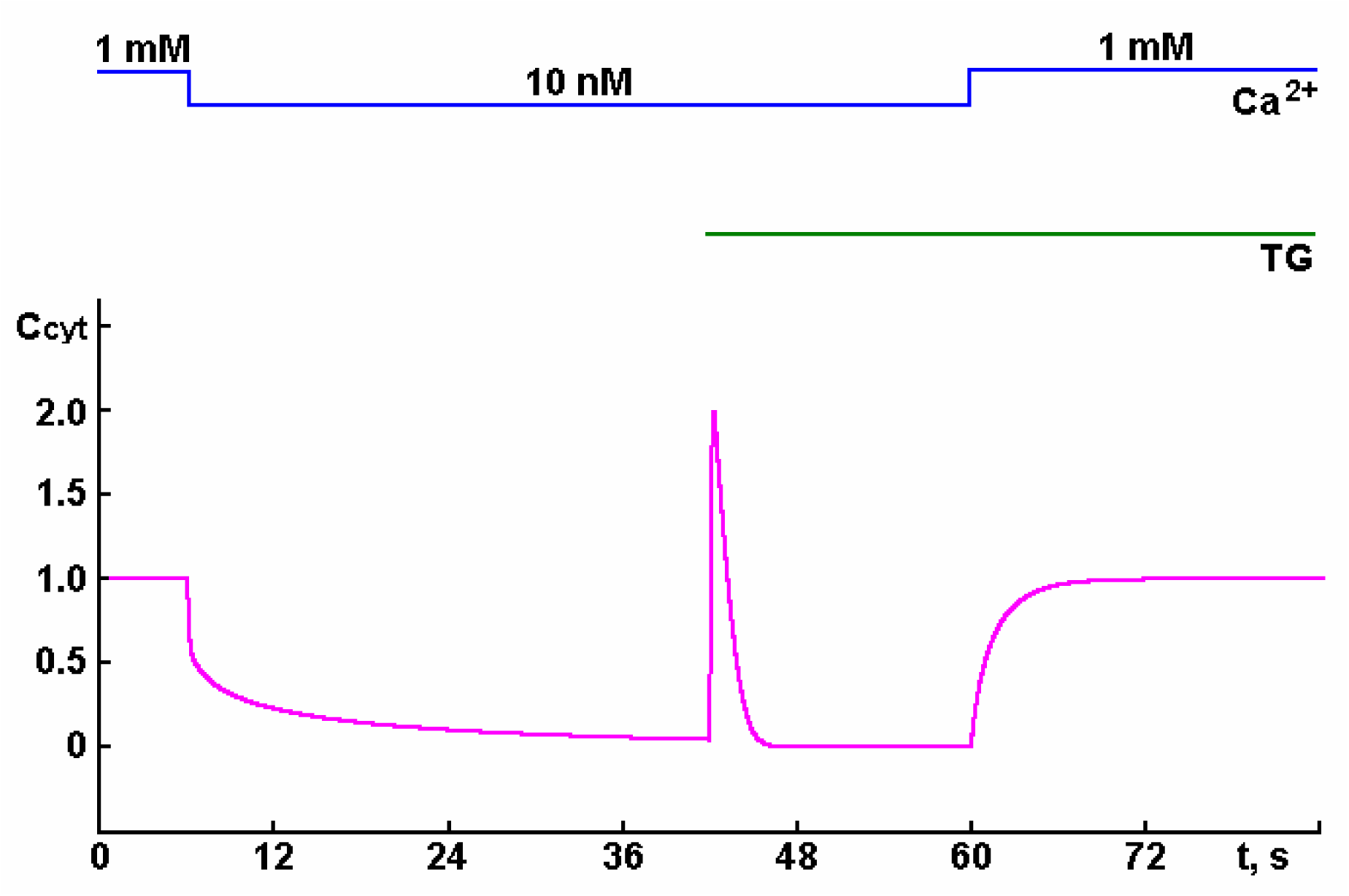
Simulation of Ca^2+^ concentration change in cytosol *C*_*cyt*_ = [*Ca*^*2+*^]_*c*_ */* [*Ca*^*2+*^]_*c,b*_ over time *t* with complete inhibition of calcium pump of SR by thapsigargin TG (*V*_*mSR*_ = 0 at 42 s) in calcium-free medium ([*Ca*^*2+*^]_*o*_ = 10 nM from 6 to 60 s). Addition of Ca^2+^ to the external medium (at a basal concentration of 1 mM) leads to a restoration of the basal Ca^2+^ concentration in the cytosol. Simulations were performed at *μ*_*SR*_ = 2000, [*A*]_0_ = 0 and [*F*]_0_ = 0. The values of other parameters are given in Table 1.

As a result, the Ca^2+^ concentration in the cytosol and SR decreases, reaching a new steady state level over time (5.57.10^-11^ M in the cytosol, 3.33.10^-6^ M in the SR, and in the extracellular solution the Ca^2+^ concentration increases up to 1.11.10^-6^ M). The reduction of *V*_*mSR*_ to zero after that (i.e., complete inhibition of the calcium pump of SR) induces a sharp increase of the cytosolic Ca^2+^ concentration with a further smooth decrease to almost the previous level (5.59.10^-11^ M). Inhibition of the calcium pump of SR also leads to a decrease of the Ca^2+^ concentration in the SR: it becomes equal to the Ca^2+^ concentration in the cytosol. As a result, the possibility of Ca^2+^ release from the SR no longer exists, since there is no difference in Ca^2+^ concentrations in the SR and in the cytosol. Therefore, when Ca^2+^ is returned to the extracellular solution (up to the basal level), the Ca^2+^ concentration slowly increases simultaneously in SR and cytosol up to the cytosolic basal level, i.e., up to 1.10^-7^ M. This increase implies Ca^2+^ entry into cytosol (and into SR) from the extracellular solution through PM and is interpreted by researchers as a proof of the presence of SOCs on the PM.

However, as the simulation shows, such an increase in cytosolic Ca^2+^ concentration may occur due to the presence of slow basal flow through the PM. Therefore, this possibility should also be taken into account when interpreting the experimental results.

This result shows that the developed model reproduces the phenomena that are usually interpreted as a manifestation of SOCE associated with the existence of SOCs on the PM. Therefore, we can conclude that the nature of the SOCE phenomenon may be related to the property of the PM to pass Ca^2+^ into the cytosol from the extracellular space due to the SBF-PM. Via such a process, SR refilling can also occur without the participation of a special class of calcium channels – store-operated calcium channels.

## 4. Conclusions

The simplest model of calcium-induced calcium release from the SR has been developed that takes into account the minimum possible number of components required for smooth muscle cell functioning:

a. two membranes (plasma membrane and sarcoplasmic reticulum membrane) with Ca^2+^ channels whose rate is proportional to the Ca^2+^ concentration gradient at both sides of the membrane, and calcium pumps that remove Ca^2+^ from the cytosol at a rate that depends on the cytosolic Ca^2+^ concentration;
b. calcium channel receptors, to which signal-transmitting substances (agonist for PM and Ca^2+^ for SR) can bind, as a result the channels opens, increasing their throughput capacity;
c. sites of Ca^2+^ binding with calcium channels on the cytosolic side of the PM responsible for inactivation of open calcium channels by cytosolic Ca^2+^;
d. slow basal flux of Ca^2+^ into the cell from the extracellular space through the PM, by which the cell system is “adjusted” so that passive and active transport through PM and SR membrane are balanced, allowing to maintain basal concentrations in cytosol, SR and extracellular space (100 nM, 1 mM and 1 mM, respectively): this flux is crucial for the model.

The developed model makes it possible to study the effect of agonist and modulation of the activity of calcium pumps of the PM and the SR on changes in the Ca^2+^ concentration in the cytosol, SR and extracellular space under both *in vivo* and *in vitro* conditions.

A study of this model, which simulates the behavior of a smooth muscle cell when it is stimulated by an agonist, leads to the following conclusions:

- the results of the simulation well agree with experimental observations;
- after the termination of stimulation, the cell system returns to the initial state in the *in vivo* mode, and under *in vitro* conditions it reaches a new stationary level;
- despite the high values of rate constants of binding of signal substance with receptors of the calcium channel of PM and cytosolic Ca^2+^ with receptors of the calcium channel of SR, the number of opened channels on the PM and membrane of SR remains rather insignificant;
- SR can act both as a passive participant in the process of Ca^2+^ accumulation in the smooth muscle cell, acting as a buffer, and play a major role in this process, significantly increasing the Ca^2+^ concentration in the cytosol due to calcium-induced calcium release, which is initiated by Ca^2+^ entry from the extracellular space through the PM;
- under certain conditions, depending on the values of the model parameters, both single cytosolic Ca^2+^ transient mode and a periodic oscillation mode can be observed. The periodic oscillation mode, as well as the single cytosolic Ca^2+^ transient mode, is characterized by changes in Ca^2+^ concentrations not only in the cytosol but also in the extracellular space and SR. The possibility of Ca^2+^ redistribution between the three compartments (extracellular space, cytosol, and SR) creates the possibility of cytosolic Ca^2+^ concentration oscillations;
- for the process of calcium-induced calcium release, there is no particular need to involve the idea that cytosolic Ca^2+^ must close the calcium channels of the PM to terminate Ca^2+^ flow from the extracellular space into cytosol;
- the termination of calcium-induced calcium release occurs spontaneously due to the fact that Ca^2+^ fluxes into the cytosol and out of the cytosol become equal: the active Ca^2+^ flux from the cytosol increases (due to an increase in the rate of calcium pumps of the PM and SR caused by an increase in the Ca^2+^ concentration in the cytosol), while the passive Ca^2+^ flux into the cytosol first increases (due to the increased number of open calcium channels on the SR membrane) and then begins to decrease (due to a decrease in the Ca^2+^ concentration gradient both between the cytosol and the extracellular space and between the cytosol and the SR); as a result, at a certain time point these fluxes become equal and the process of Ca^2+^ accumulation in the cytosol is terminated. Thus, calcium-induced calcium release is terminated for purely kinetic reasons and not because of the closure by cytosolic Ca^2+^ of calcium channels on the PM opened by the agonist. As a result of calcium-induced calcium release, the sarcoplasmic reticulum is not completely emptied, but rather significant amounts of Ca^2+^ remain in it;
- the model qualitatively reproduces the results of experimental studies performed to identify SOCs using inhibitors of calcium pump of SR in calcium-free medium;
- the process, which is interpreted as refilling of SR with Ca^2+^ from extracellular space through SOCs, may be of a purely kinetic nature: the Ca^2+^ flux from extracellular space into SR through cytosol is caused by the fact that at this time there is Ca^2+^ excess in extracellular space and Ca^2+^ deficiency in cytosol and SR (in comparison with basal level). This determines the appearance of the driving force for Ca^2+^ transfer from extracellular space to cytosol and from cytosol to SR;
- the process of SR refilling can proceed at the expense of SBF-PM that becomes noticeable when the Ca^2+^ concentration in the extracellular space exceeds the basal level, and the Ca^2+^ concentration in the SR and cytosol is below the basal level. This process is similar to SOCE, but does not need a separate type of calcium channels (SOCs). Ca^2+^ from the extracellular space is transported through the cytosol into SR, and this process takes place (due to the specific kinetics of the process) at the cytosolic Ca^2+^ concentration close to the basal level. Note that the emptying of SR after inhibition of the calcium pump of SR also occurs at the expense of slow basal calcium flow from the SR into the cytosol (SBF-SR). This does not require the opening of calcium channels on the SR membrane, but only needs to stop pumping out Ca^2+^ from the SR into the cytosol.

We understand that the simple model developed by us cannot fully coincide with the experiment. However, our study allows us to have a fresh view on the process of calcium-induced calcium release from SR and demonstrate that another interpretation of the experimental results is also possible. The ideas about the mechanism of calcium-induced calcium release from SR as well as the mechanism of SR refilling presented in this article should also be considered to be potentially possible, along with other concepts about the mechanisms providing calcium homeostasis in muscle cells.

